# Phosphoantigen-Driven Dissociation of Butyrophilin Oligomers Activates γδ T Cells

**DOI:** 10.1101/2025.09.26.677252

**Authors:** Weizhi Xin, Bangdong Huang, Weijie Gao, Wenjia Zhang, Yundi Hu, Yuehua Liu, Enyuan Liang, Yigong Shi, Qiang Su, Qiang Zhou

**Author notes:** Corresponding author. (Q.S.); (Q.Z.). These authors contributed equally: Weizhi Xin, Bangdong Huang.

## Abstract

γδ T cells represent a promising avenue for cancer immunotherapy. The Vγ9Vδ2 T cell receptor (TCR), expressed by the predominant subset of γδ T cells, responds to phosphoantigen (pAg)-engaged butyrophilins (BTNs) on various cancer cells. However, the molecular mechanism underlying pAg-mediated activation of Vγ9Vδ2 TCRs remains a subject of debate. Here, we employed an integrative approach to elucidate the mechanism of pAg reactivity in Vγ9Vδ2 T cells. Our results demonstrate that BTNs form higher-order oligomers in the absence of pAg. Upon pAg binding, these oligomers dissociate into tetramers, enabling Vγ9Vδ2 TCR engagement. This pAg-induced dissociation of higher-order BTN complexes is critical for pAg-mediated activation of γδ T cells. Our findings reveal a mechanism of BTN dissociation-driven pAg sensing, providing valuable insight for future immunotherapeutic strategies.

## Introduction

γδ T cells are a unique population of immune cells that can recognize and kill tumors^1^. Unlike αβ T cells, which recognize neoantigenic peptides presented by major histocompatibility complex (MHC) molecules on tumor cells, γδ T cells can detect a wide range of ligands in an MHC-independent manner^2–4^. This characteristic is particularly advantageous in certain malignancies where MHC expression is downregulated and the mutational burden is low, exemplified by pancreatic adenocarcinoma^5^. As a result, γδ T cells have emerged as promising candidates for cancer immunotherapy, especially for tumors with reduced MHC expression or limited neoantigen presentation^5^.

In humans, Vγ9Vδ2 T cells are the predominant circulating γδ T cell subset, accounting for 1–5% of total blood T cells^6^. During certain bacterial and parasitic infections, the population of Vγ9Vδ2 T cells can expand dramatically, reaching up to 50% of circulating T cells^7,8^. Vγ9Vδ2 T cells exhibit a range of effector functions, including lysing tumor cells, producing cytokines and chemokines, and priming αβ T cells via antigen presentation^2,3^. Owing to their distinct roles in tumor immunity, Vγ9Vδ2 T cells are promising targets for cancer immunotherapy^5^.

Vγ9Vδ2 T cells are activated by small non-peptidic phosphorylated metabolites referred to as phosphoantigens (pAgs)^9,10^. pAgs are naturally produced by microorganisms through the non-mevalonate pathway, such as (E)-4-hydroxy-3-methyl-but-2-enyl-pyrophosphate (HMBPP)^11,12^, or by human cells via the mevalonate pathway, exemplified by isopentenyl pyrophosphate (IPP)^10,12^. In the tumor microenvironment, dysregulation of the mevalonate pathway leads to IPP accumulation, which triggers Vγ9Vδ2 T cell activation^13–15^. Aminobisphosphonates, such as zoledronate, enhance the intracellular accumulation of endogenous pAgs and are used in certain clinical trials to boost γδ T cell activity in cancer patients^16,17^.

The activation of Vγ9Vδ2 T cells by pAgs depends on BTN3A1 and BTN2A1^18–20^. Notably, recent studies demonstrated the essential roles of BTN3A2 or BTN3A3 in pAg responses^21–25^. BTN3A2 or BTN3A3 is required for the cell surface expression of BTN2A1 and BTN3A1. Genetic knockout of both BTN3A2 and BTN3A3 abolishes Vγ9Vδ2 T cell activation, indicating that BTN3A2 or BTN3A3 plays an indispensable role in the recognition of the BTN complex by the Vγ9Vδ2 TCR.

BTN2A1, BTN3A1, and BTN3A3 exhibit similar structural features, comprising an IgG-like variable (IgV) domain and an IgG-like constant (IgC) domain in the extracellular region, a transmembrane (TM) domain, a juxtamembrane (JM) domain and a B30.2 domain in the intracellular region^26,27^. In contrast, BTN3A2 lacks the intracellular B30.2 domain. Previous studies have elucidated the interactions between BTN molecules and the Vγ9Vδ2 TCR, demonstrating that the IgV domain of BTN2A1 binds to the Vγ9 domain of the TCRγ chain^19,20^. Furthermore, the BTN2A1 homodimer forms a tetrameric complex with the BTN3A1/3A2 or BTN3A1/3A3 heterodimer^28,29^. Recent studies have proposed two models for pAg-mediated T cell activation^28,29^. The first model describes a “plier-like gripping” mechanism, in which pAg facilitates the opening of the BTN ectodomain, enabling Vγ9Vδ2 TCR binding and subsequent T cell activation^29^. The second model suggests that pAg promotes the dimerization of BTN2A1 and BTN3A1 to form a tetrameric complex that engages the Vγ9Vδ2 TCR and initiates activation^28^.

Currently, the molecular mechanisms underlying Vγ9Vδ2 T cell activation by pAgs remain a subject of debate. In this study, we determined a series of cryo-electron microscopy (cryo-EM) structures of full-length BTN molecules. Through biochemical, biophysical, and cellular analyses, we demonstrate that BTN molecules assemble into higher-order oligomers on the cell membrane. Notably, pAg binding triggers the dissociation of these oligomers into tetramers, thereby facilitating Vγ9Vδ2 TCR binding and subsequent activation of Vγ9Vδ2 T cells.

## Results

### BTN molecules form higher-order oligomers in the absence of pAgs

To explore the molecular organization of BTN molecules, we fused affinity tags to the C-termini of BTN2A1 and BTN3A1 and co-expressed them with BTN3A2 or BTN3A3 in mammalian cells (Figure S1A). Following tandem affinity purification, we subjected the purified complexes to size exclusion chromatography in the absence of pAgs (Figure S1B). For clarity, we designate the BTN2A1/3A1/3A2 complex as the 3A2 complex and the BTN2A1/3A1/3A3 complex as the 3A3 complex.

We analyzed the structures of the 3A2 and 3A3 complexes in the absence of pAgs using cryo-EM (Figures S1C and S1D). Both complexes can form higher-order oligomers without HMBPP (Figure 1A). These oligomers consist of multiple tetrameric 3A2 or 3A3 complexes arranged side-by-side, with tightly packed ectodomains. The intracellular regions appear disordered, indicating substantial conformational flexibility in the JM and B30.2 domains in the absence of HMBPP (Figure 1A).

**Figure 1.**
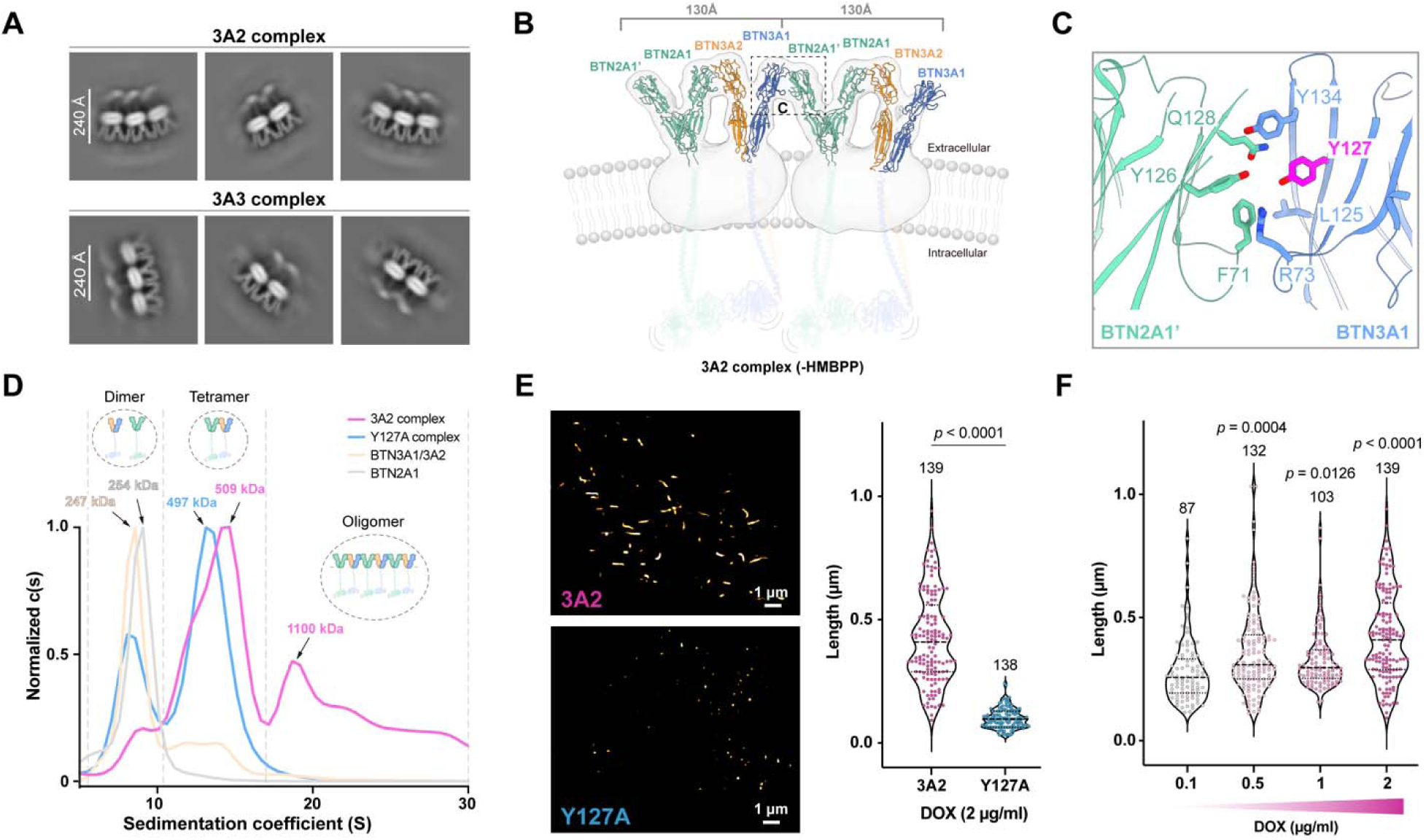
BTN molecules form higher-order oligomers on cell membrane. (A) 2D class averages demonstrate that both WT 3A2 complex (top panel) and WT 3A3 complex (bottom panel) form oligomers in the absence of HMBPP. (B) Docking of two ectodomains of the 3A2 complex into the cryo-EM maps of the WT 3A2 complex oligomer (Figure S1E). The cryo-EM map is contoured at 3σ. The ectodomains of the 3A2 complex were flexibly docked into the map using the Namdinator server. Flexible transmembrane and cytoplasmic regions (without resolvable cryo-EM density) are shown as transparent cartoons (Video S1). (C) Y127^BTN3A1^ is located in the IgV^BTN2A1^-IgV^BTN3A1^ interface (PDB ID: 8DFX^33^). (D) Analytical ultracentrifugation (AUC) reveals distinct sedimentation profiles of 3A2 complex, BTN2A1, and Y127A complex in the absence of HMBPP. Peaks at around 497-508 kDa correspond to the tetrameric assembly embedded in micelles, while the peak at ∼1,100 kDa corresponds to higher-order oligomers. Y127A, 3A2 complex variant harboring a Y127A mutation in BTN3A1. Raw data were normalized to 0-1 in GraphPad. (E) Left panel: Zoomed TIRF-SIM images of cells expressing WT 3A2 complex or Y127A complex. Right panel: Violin plots of length distribution for filaments identified in the SIM reconstruction. The WT 3A2 complex in cells induced by 2 µg/ml Dox assembles into filaments containing ∼7-85 repeats of the tetrameric unit. *P* value was determined by a Mann-Whitney test. DOX, doxycycline. **f,** Violin plots of length distribution for filaments identified in the SIM reconstruction of cells expressing WT 3A2 complex induced with different concentrations of DOX. *P* values by Kruskal-Wallis test with Dunn’s multiple comparisons post-test. Lines indicate median values.

Using our standard data processing pipeline^30–32^, we obtained a cryo-EM map of moderate resolution (Figures S1E*–*S1G). As shown in 2D averages, the intracellular regions are unresolved, suggesting substantial conformational flexibility in the JM and B30.2 domains of the oligomer (Figure 1B). Flexible docking of the ectodomain structures of the tetrameric 3A2 complex (introduced in the later text) into the map reveals a side-by-side arrangement of two 3A2 complexes, likely bridged by the IgV domains of BTN3A1 and BTN2A1’, consistent with the recently reported crystal structure^33^ (Figure 1B). Extensive van der Waals interactions are observed at the interface between IgV_BTN2A1’_ and IgV_BTN3A1_. Notably, Tyr127 has a critical role in this interface, as the Y127A mutation abolishes the interaction between BTN2A1 and BTN3A1^33^ (Figure 1C).

To investigate whether BTN molecules assemble into higher-order oligomers in the absence of pAgs, we assessed their oligomerization state in solution by analytical ultracentrifugation (AUC). Purified BTN2A1 and BTN3A1/BTN3A2 displayed peaks corresponding to molecular weights of 254 kDa and 247 kDa, respectively (Figure 1D), consistent with homodimeric and heterodimeric configurations. The purified 3A2 complex exhibited a peak at 509 kDa, matching the expected tetrameric assembly. However, additional higher molecular weight species (HMWS), including a prominent peak at approximately 1,100 kDa (Figure 1D), were also detected, suggesting the existence of oligomers larger than the tetrameric form. Notably, the Y127A mutation in BTN3A1 (referred to as Y127A), which is predicted to interfere with higher-order oligomerization (Figures 1B and 1C), selectively eliminated the HMWS while maintaining the tetrameric peak (Figure 1D). These findings provide biochemical evidence supporting the BTN higher-order oligomerization without pAgs.

To further confirm the formation of higher-order BTN oligomers on the cell membrane, we performed super-resolution imaging using total internal reflection fluorescence structured illumination microscopy (TIRF-SIM) in NIH-3T3 cells expressing either wild-type (WT) or mutant 3A2 complex variants (Figure 1E, left). TIRF-SIM imaging reveals that the WT 3A2 complex forms linear, filamentous clusters on the cell membrane, with lengths ranging from ∼100 to 1,000 nm. In contrast, the Y127A complex forms small, homogeneous puncta with significantly shorter clusters, in which the median length is less than 100 nm (Figure 1E, right). Given that each tetrameric unit spans approximately 130 Å, the observed clusters correspond to ordered arrays of 8–80 repeating tetrameric 3A2 complex units. These results demonstrate that the overexpressed 3A2 complex assembles into extended, hundreds of nanometer-scale filaments on the cell membrane, and that the IgV_BTN3A1_–IgV_BTN2A1’_ interface is critical for this oligomerization.

To determine whether 3A2 complex oligomerization is concentration-dependent, we generated cell lines with inducible and tunable expression of the 3A2 complex. Reducing the concentration of doxycycline led to a slight decrease in the median length of the filamentous clusters (Figure 1F). However, the predominant population of filamentous clusters remained above 250 nm in length, indicating that even at low expression levels, the 3A2 complex retains the capacity to form higher-order oligomers on the cell membrane. This analysis shows that filament formation is relatively consistent across varying expression levels and appears to represent an intrinsic property of BTN molecules on the cell membrane.

### pAgs drive the dissociation of higher-order BTN oligomers

To investigate the role of pAgs on BTN molecules, we successfully obtained cryo-EM structures of the 3A2 complex and 3A3 complex in complex with HMBPP at overall resolutions of 3.0 Å and 3.8 Å, respectively (Figures S2 and S3). In the presence of HMBPP, cryo-EM structures show that both 3A2 and 3A3 complexes adopt tetrameric architectures, each composed of a BTN2A1 homodimer paired with either a BTN3A1/3A2 or BTN3A1/3A3 heterodimer, arranged in a scissor-like conformation as previously reported^28,29^ (Figures 2A and S4A). The assembly mechanism of the pAg-bound 3A2 complex and the 3A3 complex is the same. The following structural analysis takes the 3A2 complex as an example. Structural analysis identifies three critical interfaces required for tetramer formation: (1) IgV–IgV domain interactions between BTN2A1 and BTN3A2; (2) extensive contacts within the TM and JM regions of the BTN2A1 homodimer, the BTN3A1/BTN3A2 heterodimer; (3) B30.2–B30.2 domain interactions between BTN2A1 and BTN3A1, glued by HMBPP (Figures S4B*–*S4F). These interfaces are involved in a network of disulfide bonds, hydrogen bonds, cation-π interactions, and van der Waals forces (Figure S5).

**Figure 2.**
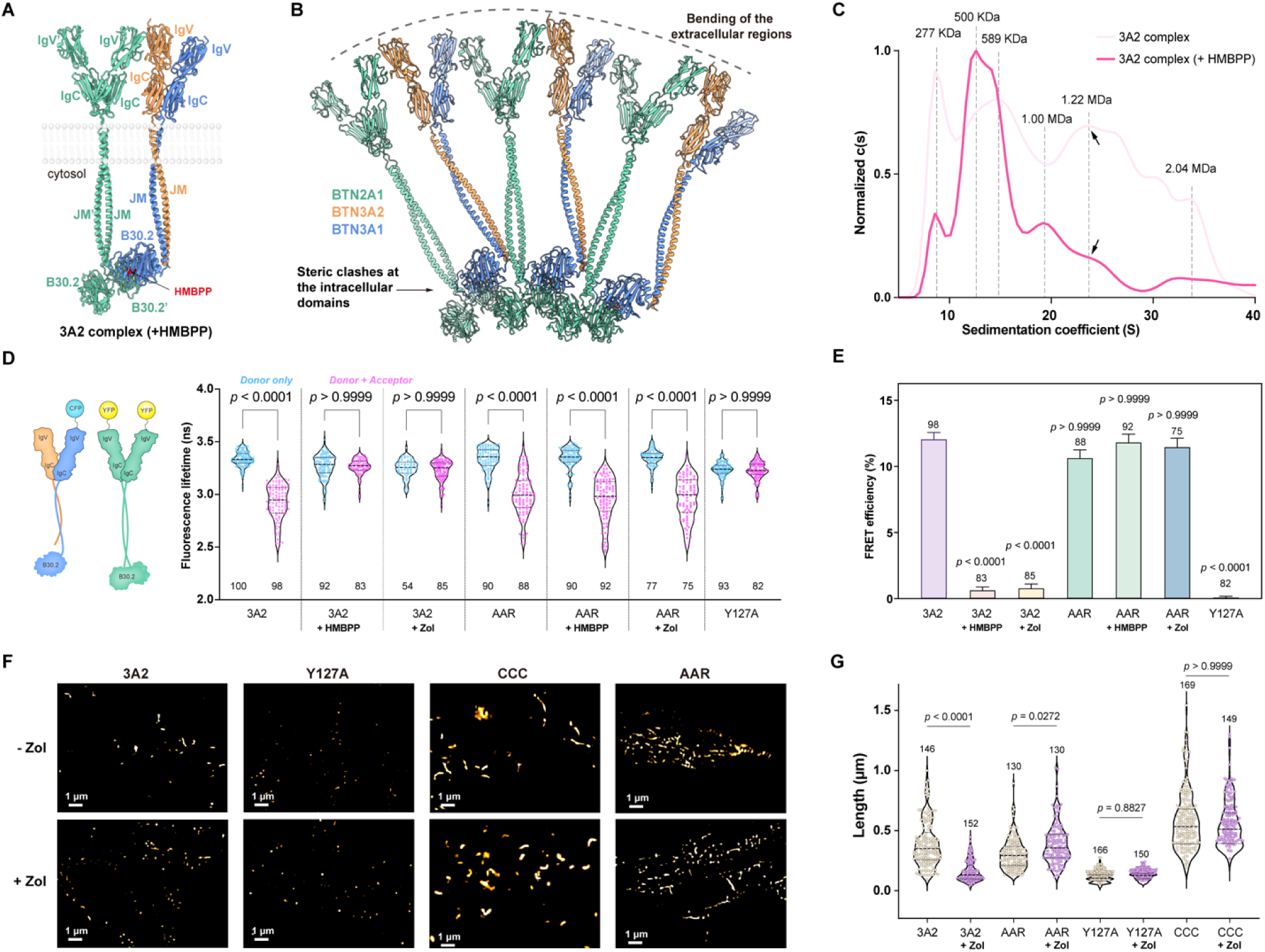
pAgs induce the dissociation of BTN higher-order oligomers. (A) The full-length structure of the HMBPP-bound 3A2 complex. The complex consists of a BTN2A1 homodimer and a BTN3A1/BTN3A2 heterodimer (Figure 4A, left). BTN2A1 interacts with BTN3A2 via IgV domains on the extracellular side and with BTN3A1 via B30.2 domains on the intracellular side. HMBPP is shown in sphere style in red. See also Figures S4B–S4F for more detailed interaction information. The statistics for the model are shown in Table S1. (B) Multiple HMBPP-bound 3A2 complexes were concatenated based on the IgV_BTN2A1_– IgV_BTN3A1_ interface (PDB ID: 8DFX^33^). Binding of HMBPP induces the bending of the extracellular region of the BTN complex and causes spatial conflicts at the intracellular domains, which collectively destabilize the higher-order oligomeric assembly. (C) Analytical ultracentrifugation (AUC) sedimentation profiles of the WT 3A2 complex (± 40 μM HMBPP). HMBPP drives the dissociation of high-mass oligomers into lower-mass species. Raw data were normalized to 0-1 in GraphPad with two aberrant points removed. (D) FLIM-FRET assays assessing the interaction between BTN2A1 and BTN3A1. Cartoons depict the construct design for the FRET reporter construct. The fluorescence lifetime of CFP was measured in Lenti-X 293T cells. “Donor only”: Cells expressing BTN3A1 with N-terminally fused CFP. “Donor + Acceptor”: Cells co-expressing BTN3A1 with N-terminally fused CFP and BTN2A1 with N-terminally fused YFP (Figures S7A and S7B). The N value is provided for each group (mean ± SEM). *P* values were calculated by the Kruskal-Wallis test with Dunn’s multiple comparisons post-test. Zol, zoledronate. Y127A, the 3A2 complex variant harboring a BTN3A1-Y127A mutation to disrupt the oligomeric assembly; AAR, the 3A2 complex variant harboring BTN3A1-H381R/BTN2A1-R477A/T510A mutations to abolish HMBPP binding. (E) FRET efficiency was determined in cells expressing the 3A2 complex, AAR, and Y127A variants. The mean fluorescence lifetime in each “Donor only” group was used as a control to calculate FRET efficiency in each “Donor + Acceptor” group. The N value is provided for each group (mean ± SEM). *P* values were calculated using the Kruskal-Wallis test with Dunn’s multiple comparisons post-test. (F) TIRF-SIM super-resolution imaging of NIH-3T3 cells expressing the WT 3A2 complex or its variants (± 10 μM Zol). Raw images were cropped to highlight regions of interest. CCC, the 3A2 complex variant harboring G130C in BTN2A1, D132C in BTN3A1, and D132C mutation in BTN3A2, respectively, to lock the oligomeric state by intermolecular disulfide bonds. (G) Violin plots of length distribution for all filaments identified in the SIM images from (F). *P* values were calculated using the Kruskal-Wallis test with Dunn’s multiple comparisons post-test.

Compared to the crystal structure of the BTN2A1 and BTN3A1 extracellular domains complex, which can assemble into high-order oligomers in the crystal (Figure S6A), the extracellular domains of the HMBPP-bound 3A2 complex exhibit substantial conformational changes. The interchain angle between the extracellular domains of BTN2A1 and BTN3A2 in HMBPP-bound 3A2 complex is decreased by 7.1° (Figure S6B), while the dimerization angles of the extracellular domains are increased by 3.5° for the BTN2A1 dimer and by 7.7° for the BTN3A1/BTN3A2 heterodimer, respectively (Figures S6C and S6D). The assembly of the HMBPP-bound 3A2 tetrameric complexes into the oligomer based on the IgV_BTN2A1_–IgV_BTN3A1_ interface in the crystal structure of the BTN2A1 and BTN3A1 extracellular domains complex (PDB ID: 8DFX) reveals curvature at the extracellular regions and steric clashes at the intracellular domains of BTN molecules (Figure 2B). In the absence of pAgs, the JM and B30.2 domains are flexible. Upon HMBPP binding, these domains are stabilized by HMBPP, giving rise to steric hindrance at the intracellular region, accompanying curvature change at the extracellular regions induced by the conformational changes at the extracellular regions of the BTN complex, thus jointly destabilizing the higher-order oligomer of 3A2 complex and triggering the dissociation of higher-order BTN oligomers into discrete tetrameric 3A2 complex (Video S1).

To test the hypothesis that pAg induces the dissociation of oligomers, we first assessed the effect of pAg on the oligomerization state of the 3A2 complex in solution using AUC (Figure 2C). Compared to the WT 3A2 complex in the absence of HMBPP, the proportion of HMWS, corresponding to the molecular weights exceeding 500 kDa, was significantly reduced upon HMBPP addition. These results suggest that HMBPP induces the dissociation of higher-order BTN oligomers into tetrameric units of 3A2 complex.

To investigate the oligomerization states of BTN molecules on cell membrane, we employed fluorescence lifetime imaging microscopy combined with fluorescence resonance energy transfer (FLIM-FRET). CFP and YFP were fused to the N-termini of BTN3A1 and BTN2A1, respectively, and co-expressed with BTN3A2 (Figures 2D, S7A, and S7B). Cells expressing the WT 3A2 complex exhibited a marked reduction in CFP fluorescence lifetime, indicating close proximity among those tetrameric 3A2 units and suggesting higher-order oligomerization. In contrast, treatment with HMBPP or zoledronate abolished the change in CFP fluorescence lifetime, indicating dissociation of the higher-order BTN oligomers (Figures 2D and 2E). As controls, we generated a 3A2 complex variant carrying the H381R mutation in BTN3A1 and the R477A/T510A mutations in BTN2A1 (referred to as AAR) to disrupt pAg binding^34^, or the Y127A mutation in BTN3A1 to disrupt higher-order oligomer formation (Figure 1E). As expected, the FRET signal of the Y127A complex was diminished, and the FRET signal of the AAR complex remained unchanged upon pAg treatment (Figures 2D and 2E).

To further validate these findings, we used an alternative approach by fusing CFP or YFP to the C-terminus of BTN3A2 (Figures S7C and S7D). Consistent with previous results, cells expressing the WT 3A2 complex exhibited a strong FRET signal, which was eliminated by pAg treatment, supporting that pAg induces dissociation of BTN higher-order oligomers. In control experiments, the FRET signal was absent in cells expressing the Y127A complex, and the AAR complex remained unaffected by pAg treatment (Figures S7E and S7F). Collectively, these FRET data show that the 3A2 complex forms higher-order oligomers on the cell membrane, and that this oligomerization is disrupted upon pAg treatment.

To substantiate the model that pAg induces the dissociation of BTN oligomers, we employed TIRF-SIM to monitor alterations in BTN clusters on the cell membrane following pAg treatment. For WT 3A2 complex, pAg addition led to a significant decrease in the length of membrane-associated clusters (Figures 2F and 2G). However, when BTN oligomers were locked by intermolecular disulfide bonds (CCC mutant) or pAg sensing was disrupted by AAR mutations, pAg was unable to induce cluster dissociation. Consistent with our model, pAg treatment did not affect the length of membrane-bound clusters in cells expressing the Y127A complex, in which higher-order oligomer formation is already compromised (Figures 2F and 2G).

In summary, we demonstrate that HMBPP binding triggers the disassembly of BTN higher-order oligomers. We propose that pAgs binding induces curvature changes in the extracellular regions and rigidifies the intracellular segments of BTN molecules, leading to steric clashes at the intracellular domains. To corroborate this model, we used AUC to monitor the dissociation of BTN oligomers in solution upon pAg binding. Additionally, we evaluated oligomer disassembly on the cell membrane using FLIM-FRET and TIRF-SIM. Collectively, these findings show that pAg induces the dissociation of BTN higher-order oligomers, revealing a mechanism distinct from recently proposed models^28,29^.

### Vγ9Vδ2 TCR binds to BTN tetramer independently of pAg

The 3A2 or 3A3 complex can associate with either the G115 TCR^9^ or MOP TCR^35,36^, in the presence of HMBPP, as shown by gel filtration analysis (Figures S8A–S8C). We determined the structures of the G115 TCR–3A2, MOP TCR–3A2, and G115 TCR–3A3 complexes at resolutions ranging from 3.3 to 3.8 Å in the presence of HMBPP (Figures S9 and S10). These structural findings, consistent with recent studies^28,29^, reveal a widely accepted dual-ligand recognition mechanism. The G115 TCR binds to BTN ectodomains, with the Vγ9 chain interacting with IgV_BTN2A1_ laterally, while the Vδ2 chain apical face engages IgV_BTN3A2_ (Figures S11A and S11B). A similar interaction pattern is observed in the MOP TCR–3A2 complex (Figure S11C). Interestingly, across all datasets of these structures, we consistently observed a small subset of particles in which another TCR binds exclusively to the BTN2A1 IgV domain, consistent with the recent report^28^ (Figure S12).

The G115 TCR–3A2 complex reveals that the recognition of the BTN complex by Vγ9Vδ2 TCR involves both somatically recombined complementarity-determining region 3 (CDR3) and germline-encoded regions such as CDR1, CDR2, and hypervariable 4 (HV4) (Figure S11D). This recognition mode is consistent with prior functional studies showing the necessity of both CDRs and HV4 regions for BTN recognition by Vγ9Vδ2 TCR^19,20,37,38^. Structural analysis delineates intricate interaction networks of IgV_BTN3A2_–Vδ2 and IgV_BTN2A1_– Vγ9 interfaces involving at least 18 hydrogen bonds, two cation-π interactions, and extensive van der Waals forces (Figures S11E and S13).

Our structures also provide a structural framework explaining decades of mutagenesis data related to pAg-induced activation of Vγ9Vδ2 TCR^19,20,22,33,34,37–40^. Mapping these mutations onto the G115 TCR–3A2 structure shows that all those previously identified critical mutations localize precisely to the interfaces between the G115 TCR and 3A2 complex, as observed in our structures (Figure S11F). The striking concordance between structural and functional data underscores the physiological importance of our structures.

To investigate the role of pAg in BTN complex recognition by the Vγ9Vδ2 TCR, we determined the cryo-EM structure of the G115 TCR–AAR complex at 3.2 Å resolution (Figures S8F and S14). The AAR complex is a 3A2 complex variant carrying the H381R mutation in BTN3A1 and the R477A/T510A mutations in BTN2A1 to disrupt pAg binding^34^. Compared to the HMBPP-bound G115 TCR–3A2 structure, the extracellular domains remain well-resolved in the absence of HMBPP. However, the TMs, JMs, and B30.2 domains are invisible in the final reconstruction, indicating a considerable flexibility in these regions (Figure 3A). Structural superposition reveals that the overall extracellular architecture is similar, with a root mean standard deviation (RMSD) of 0.3 Å, indicating that the TCR–BTN interface remains unchanged.

**Figure 3.**
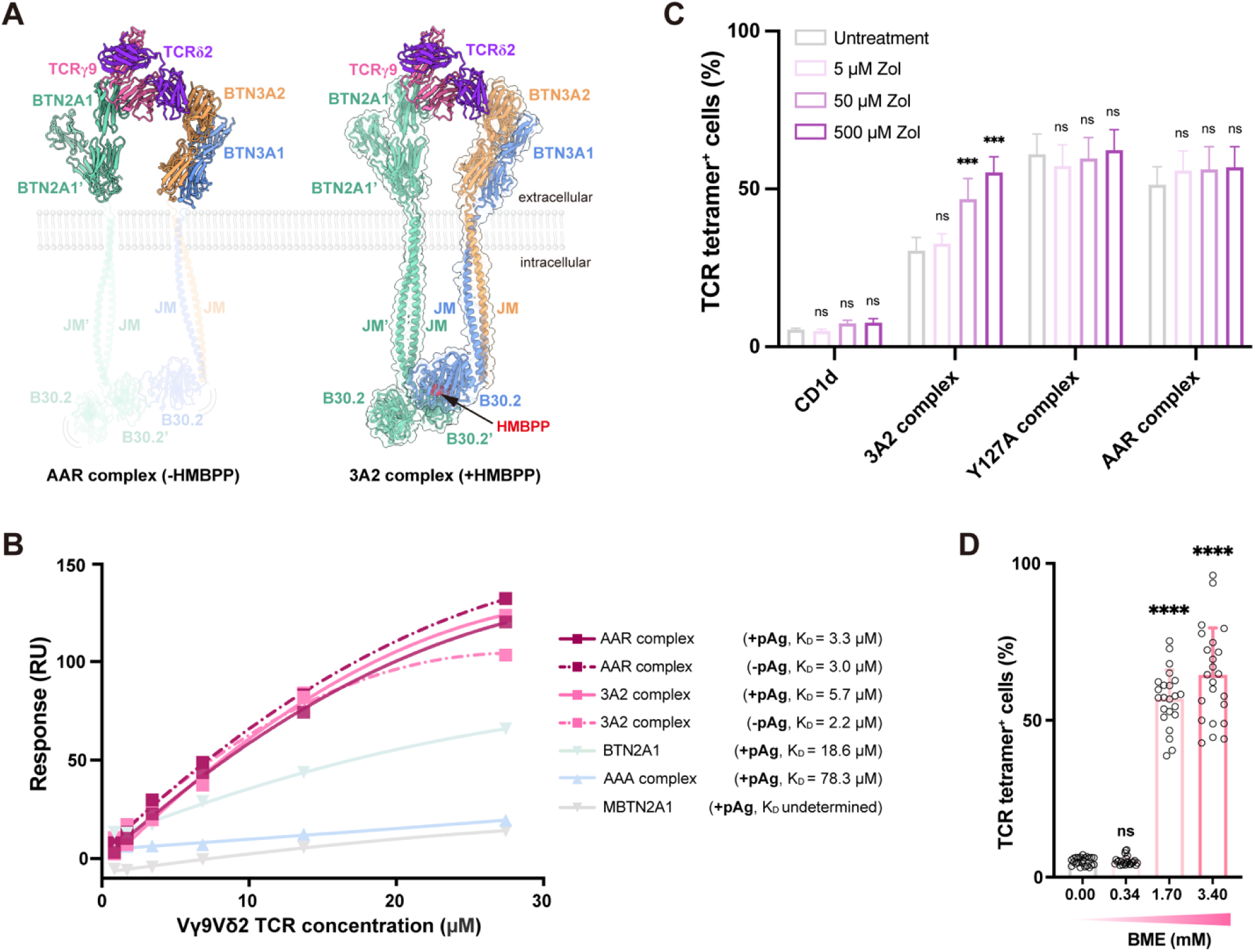
Vγ9Vδ2 TCR binding to the BTN tetramer is independent of pAgs. (A) Left: Cryo-EM structure of Vγ9Vδ2 TCR (G115 prototype) in complex with the ectodomain of AAR complex in the absence of HMBPP. The variable domains of the TCR bind IgV_BTN2A1_ and IgV_BTN3A2_. Intracellular regions exhibit conformational flexibility (unresolved in cryo-EM map) and are depicted as transparent cartoons. AAR complex, a 3A2 complex variant with mutations H381R in BTN3A1 and R477A/T510A in BTN2A1 B30.2 domains to disrupt pAg binding. Right: Overall cryo-EM structure of Vγ9Vδ2 TCR (G115 prototype) in complex with the HMBPP-bound WT 3A2 complex. The structure is presented in a cartoon style, with HMBPP highlighted in sphere style in red. BTN molecules are displayed as transparent surfaces. Subunit rendering: BTN2A1 (medium aquamarine), BTN3A1 (cornflower blue), BTN3A2 (sandy brown), BTN3A3 (khaki), TCRδ (blue violet), TCRγ (hot pink). See also Figures S11E for more detailed inter-chains interaction information. The statistics for the model are shown in Table S1. (B) Saturation binding curves of G115 TCR measured by surface plasmon resonance (SPR). WT 3A2 and AAR complexes were tested with 4 μM HMBPP or without HMBPP. WT or mutant BTN2A1, and the AAA complex were tested in the presence of 4 μM HMBPP only. AAA complex, a 3A2 complex variant carrying the R124A/Y133A mutations in BTN2A1 and D132A in BTN3A2. MBTN2A1, a BTN2A1 variant carrying E63A/R65A/F71A/K79A/R84A/E87A/R131A/E135A mutations in BTN2A1 to disrupt TCR binding. RU: resonance units. K_D_, dissociation constant at equilibrium ± SEM. (C) G115 TCR tetramer staining of cells expressing WT 3A2 complex or its variants after incubation with Zol. CD1d-expressing cells served as negative controls. The frequency of TCR tetramer+ cells was analyzed by flow cytometry. \*\*\**p* < 0.001, by two-way ANOVA with Dunnett’s multiple comparison test. Zol, zoledronate. (D) G115 TCR tetramer staining of NIH-3T3 fibroblasts transfected with the CCC complex, after incubation of the cells with BME at the indicated concentrations in the absence of pAgs. The frequency of TCR tetramer^+^ cells was analyzed by flow cytometry. \*\*\*\**p* < 0.0001, by the Kruskal-Wallis test with Dunn’s multiple comparisons post-test. BME, β-Mercaptoethanol.

Our structural data suggest that pAg does not affect the interaction between the Vγ9Vδ2 TCR and the BTN complex. To test this hypothesis, we measured binding affinities by surface plasmon resonance (SPR) (Figures 3B and S15). Purified tetrameric WT and AAR complexes showed similar binding affinities for G115 TCR in the presence of pAg, with dissociation constants (K_D_) of 5.7 μM and 3.3 μM, respectively. In the absence of pAg, the binding affinity was also similar, with K_D_ values of 2.2 μM and 3.0 μM for the respective complexes. Although cryo-EM analysis has revealed the intricate interaction network between the TCRδ chain and the BTN3A2 IgV domain, SPR analysis did not detect a measurable binding affinity between the TCR and the BTN3A1/3A2 complex, consistent with previous finding^41^. The BTN2A1 dimer showed weaker binding, with K_D_ values of 18.6 μM, comparable to those reported previously^19,20^. In contrast, the AAA complex and the MBTN2A1 dimer mutants designed to disrupt TCR binding showed markedly reduced affinities, with K_D_ values of 78.3 μM for AAA complex, while no detectable affinity for MBTN2A1 (Figures 3B and S15). These results demonstrate that, similar to the tetrameric 3A2 complex, the G115 TCR interacts with the ectodomains of BTN2A1 and BTN3A2 independently of HMBPP binding to the B30.2 domain.

### pAg drives Vγ9Vδ2 TCR binding to BTN molecules on the cell membrane

To further elucidate the role of pAgs, we performed Vγ9Vδ2 TCR tetramer staining assays to evaluate how pAgs affect Vγ9Vδ2 TCR binding to BTN molecules on the cell membrane. In cells expressing the 3A2 complex, substantial G115 TCR staining was detected in the absence of zoledronate treatment. Increasing concentrations of zoledronate markedly enhanced G115 TCR binding in these cells (Figure 3C). In contrast, cells expressing the AAR complex, which is impaired in pAg binding, showed no enhancement in G115 TCR staining following zoledronate treatment. Strikingly, cells expressing the Y127A complex, which forms a constitutive tetramer, displayed high levels of G115 TCR binding that were not further increased by zoledronate (Figure 3C). These results suggest that the dissociation of the oligomeric 3A2 complex into tetrameric units is critical for augmenting TCR binding on the cell membrane.

To verify that oligomer dissociation is necessary for pAg-induced TCR binding, we examined Vγ9Vδ2 TCR tetramer staining in cells expressing the CCC variant, which incorporates the G130C, D132C, and D132C mutations in BTN2A1, BTN3A1, and BTN3A2, respectively. These mutations were designed to stabilize the oligomeric state through the formation of intermolecular disulfide bonds. In the absence of β-mercaptoethanol (BME), Vγ9Vδ2 TCR tetramers failed to stain cells expressing the CCC complex (Figure 3D). However, treatment with BME concentrations of 1.70 mM or higher induced a marked increase in Vγ9Vδ2 TCR tetramer binding. These data confirm that pAg-induced TCR binding on the cell membrane necessitates the dissociation of 3A2 complex oligomers (Figure 3D).

In summary, these findings demonstrate that the tetrameric BTN complex directly interacts with the TCR. Notably, pAg does not affect TCR binding affinity in this tetrameric configuration. In contrast, higher-order BTN oligomers inhibit TCR binding. pAgs promote BTN–TCR interaction on cell membrane by dissociating higher-order oligomers into tetrameric units, representing a mechanism distinct from those recently reported^28,29^.

### BTN oligomer dissociation contributes to the pAg-dependent γδ T cell activation

Combining AUC, TIRF, FLIM-FRET, and tetramer staining analyses, we demonstrated that pAg induces the dissociation of high-order BTN oligomers into tetramers. To determine whether dissociation of BTN oligomers is required for pAg-mediated γδ T-cell activation, we assessed the ability of cells expressing the 3A2, Y127A, or AAR BTN complexes to activate T cells in the presence or absence of pAg. The 3A2 complex forms oligomers that dissociate into tetramers upon exposure to pAg; the AAR complex fails to bind pAg and remains as a higher-order oligomer; and the Y127A complex exists constitutively as tetramers, independent of pAg treatment.

We co-cultured primary Vγ9Vδ2 T cells, expanded from peripheral blood mononuclear cells (PBMCs), with NIH-3T3 cells expressing the 3A2, Y127A, or AAR complex, all of which exhibited comparable surface expression levels (Figures 3B and S16). Consistent with previous findings^19^, cells expressing BTN2A1 or BTN3A1/BTN3A2 alone failed to induce CD25 expression in primary Vγ9Vδ2 T cells, regardless of pAg stimulation. However, cells expressing the 3A2 complex induced substantial CD25 expression in T cells, and exposure to zoledronate further enhanced CD25 expression, demonstrating robust pAg-dependent activation (Figure 4A). Co-culturing primary Vγ9Vδ2 T cells with cells expressing the pAg-binding-defective AAR complex resulted in CD25 expression levels comparable to those induced by the 3A2 complex (Figure 4A). As recently reported^33^, the pAg-dependent augmentation of CD25 expression was reduced in this condition (Figure 4B).

**Figure 4.**
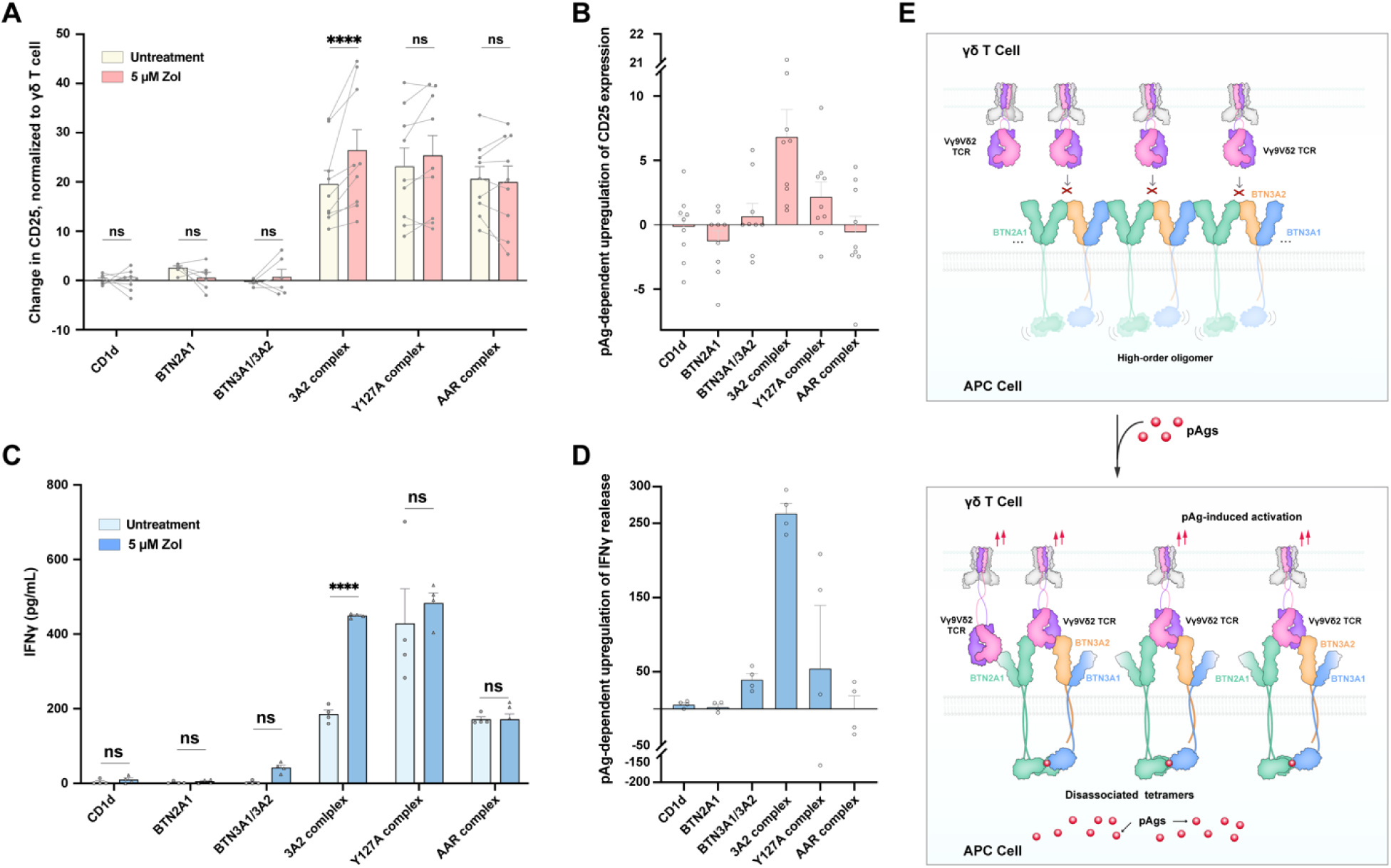
PAg responses of Vγ9Vδ2 T cell is mediated by the dissociation of BTN higher-order oligomers. (A and B) CD25 expression level in Vγ9Vδ2 T cells co-cultured with NIH-3T3 fibroblasts expressing different BTN molecules pretreated with or without 5 μM Zol (A) and upregulation level of CD25 in Vγ9Vδ2 T cells with zoledronate treatment compared with the untreated group (B). CD1d-expressing cells served as negative controls. Zol, zoledronate. \*\*\*\**p* < 0.0001, by two-way ANOVA with Sidak’s multiple correction test in (A). (C and D) IFN-γ concentration in the Vγ9Vδ2 T cell culture medium (C) and changes in IFN-γ concentration in the culture medium of Vγ9Vδ2 T cells before and after Zol treatment (D). \*\*\*\**p* < 0.0001, by two-way ANOVA with Sidak’s multiple correction test in (C). (E) Proposed model of pAg-mediated activation of Vγ9Vδ2 TCR. In the absence of pAgs, BTN molecules form oligomers on the cell membrane with flexible intracellular regions. Upon pAg binding, bending of extracellular regions and steric hindrance within the intracellular rigid segments of BTN molecules, collectively trigger the dissociation of BTN oligomers into tetramers. The Vγ9Vδ2 TCR then engages the ectodomains of the tetrameric BTN complex.

Co-culturing primary Vγ9Vδ2 T cells with cells expressing the Y127A complex also induced strong CD25 expression. Notably, the pAg-dependent increase in CD25 expression was absent in T cells co-cultured with the Y127A complex (Figure 4B), indicating that the Y127A mutation in BTN3A1 disrupts pAg-dependent activation. Importantly, in the absence of pAg, γδ T cells co-cultured with cells expressing the Y127A complex exhibited higher CD25 expression than those co-cultured with the 3A2 complex, highlighting the role of the tetrameric assembly of the BTN complex in γδ T cell activation (Figure 4B).

Similar patterns were observed in the detection of IFNγ release when primary Vγ9Vδ2 T cells were co-cultured with NIH-3T3 cells expressing the 3A2, Y127A, or AAR complex (Figures 4C and 4D). pAg-dependent IFNγ release was detected in T cells co-cultured with cells expressing the 3A2 complex, whereas IFNγ release was significantly reduced in T cells co-cultured with cells expressing the Y127A or AAR complex, consistent with the CD25 expression patterns.

In summary, based on the distinct oligomeric states of the 3A2, AAR, and Y127A BTN complexes in the presence or absence of pAg, these findings suggest that the dissociation of higher-order oligomeric BTN complexes into tetrameric units following pAg binding is essential for pAg-mediated activation of Vγ9Vδ2 T cells.

## Discussion

Our biophysical, biochemical, and cellular analyses support an unprecedented pAg-dependent γδ T cell activation model, which is distinct from those recently reported ^28,29^. In this model, BTN molecules form higher-order oligomers on the cell membrane in the absence of pAgs. Upon intracellular binding of pAgs, these oligomers dissociate into tetramers, thereby enabling γδ TCR engagement (Figure 4E).

In the absence of pAgs, BTN molecules assemble into higher-order oligomers on the cell membrane. Within these oligomers, the ectodomains adopt a tightly packed, side-by-side arrangement, whereas the cytoplasmic regions remain flexible to avoid steric hindrance. This compact ectodomain organization occludes the TCR–BTN binding interface, thereby preventing γδ TCR engagement (Figure 4E).

Binding of pAg induces curvature of the extracellular regions and steric clashes at the intracellular domains of the BTN oligomer. In the intracellular regions, pAg functions as a molecular glue, enhancing the interaction between the B30.2 domains of BTN2A1 and BTN3A1 within the higher-order BTN oligomer. This interaction stabilizes the transmembrane and juxtamembrane regions, leading to steric clashes between adjacent BTN molecules accompanying the bending of the extracellular regions of the BTN oligomer, thus liberating discrete BTN tetramers. Once uncaged, these tetramers can engage Vγ9Vδ2 TCR in a dual-ligand manner, adopting a ‘scissor-like’ conformation as previously reported^28,29^, thereby initiating γδ T cell activation (Figure 4E).

Our model is consistent with previously published data. Prior studies have shown that the E35A mutation on BTN2A1^28,29^, as well as the R73A or Y127A mutations on BTN3A1^39^, which disrupt BTN2A1–BTN3A1 interactions, significantly enhance the stimulation of Vγ9Vδ2 T cells. Among these, the Y127A or R73A mutations on BTN3A1 show the strongest increase in HMBPP-induced TCR activation^39^. This paradoxical enhancement in activation can be attributed to the destabilization and/or dissociation of the oligomer caused by these mutations. In contrast, stabilizing the conformation of oligomers or the internal BTN3A molecules disfavors dissociation of the oligomer, thereby antagonizing the pAg-induced response^33,42^.

One of the recently reported models suggests that pAg facilitates the formation of a tetrameric complex by binding a BTN2A1 dimer with either a BTN3A heterodimer or a BTN3A1 homodimer, enabling dual-ligand recognition by the Vγ9Vδ2 TCR^28^. Another model proposes that pAg stabilizes the intracellular domains of a tetrameric BTN complex into a “plier-like gripping” conformation, thereby priming it for TCR engagement^29^. However, our biochemical and cellular data show different scenarios. First, the 3A2 complex is capable of spontaneously forming high-order oligomers in the absence of pAg. Second, the Y127A complex, a constitutively stable tetramer that elicits comparable levels of CD25 upregulation and IFN-γ secretion in Vγ9Vδ2 T cells, independent of pAg presence. These findings are not reconcilable with the existing models. Nonetheless, our structural framework does not preclude an alternative role for pAg when BTN molecules are sparsely expressed on the cell surface and exist as dimeric species. pAg may act as “molecular glue”, bridging BTN2A1 and BTN3A dimers to form tetramers and induce a response^28^.

In conclusion, our research observes the oligomerization of BTN molecules in the absence of pAg and uncovers a pAg-dependent activation mechanism, whereby BTN oligomers dissociate into tetramers upon pAg stimulation, leading to the activation of Vγ9Vδ2 T cells. In contrast, in the absence of pAg, BTN molecules form oligomers that exclude the TCR, thereby preventing γδ T cell autoactivation. These structural and functional insights not only reshape our understanding of γδ T cell activation but also offer a framework for the development of BTN-targeted immunotherapies.

## Materials and methods

### Construct design

The cDNAs encoding human BTN2A1, BTN3A1, BTN3A2, and BTN3A3 were codon-optimized and synthesized by Tsingke Biotechnology Co., Ltd. These cDNAs were subsequently subcloned into an optimized pCAG vector. A synthetic signal peptide (MDMRVPAQLLGLLLLWLSGARC)^43^ was fused to the N-terminus of BTN2A1, BTN3A1, BTN3A2, and BTN3A3. To facilitate protein purification, a Flag tag (MDYKDDDDK) was inserted at the C-terminus of BTN2A1 or BTN3A3, while a twin-strep II tag (WSHPQFEKGGGSGGGSGGSAWSHPQFEK) was added to the C-terminus of BTN3A1. The coding sequences for the extracellular domains of the G115^9^ and MOP^35,36^ TCRδ2 and TCRγ9 chains were also codon-optimized and fused with the same synthetic signal peptide at their N-termini. Additionally, a 6×His tag and a cFos tag^44^ (ASTDTLQAETDQLEDEKYALQTEIANLLKEKEKLGAP) were inserted at the C-terminus of TCRδ2. The JunW_ph1_ tag^44^ (ASAAELEERVKTLKAEIYELRSKANMLREQIAQLGAP) was fused at the C-terminus of TCRγ9. Detailed information regarding these constructs can be found in the Supplementary Information.

### Protein purification of 3A2 complex and 3A3 complex

To purify the 3A2 complex, 2.4 mg of plasmids encoding BTN2A1, BTN3A1, and BTN3A2 (0.8 mg per plasmid) were pre-incubated with 5 mg of 40K polyethyleneimine (PEI) (Yeasen) in 40 mL of fresh medium (Sino Biological Inc.) for 30 minutes, as previously described^45–47^. The plasmid-PEI mixture was then transfected into 0.8 liters of ExpiHEK293F cells (Thermo Fisher Scientific Inc.). The transfected cells were cultured for 3 days at 37°C with 5% CO_2_ in a Multitron-Pro shaker (Infors, 120 rpm) before collection. The harvested cells were resuspended in a lysis buffer containing 25 mM HEPES (pH 7.4), 150 mM NaCl, 2 μg/mL pepstatin, 1 μg/mL aprotinin, and 2 μg/mL leupeptin. The cell membranes were solubilized overnight at 4°C using 1% (w/v) LMNG (lauryl maltose neopentyl glycol, Anatrace) and 0.1% CHS (cholesteryl hemisuccinate tris salt, Anatrace). The cell lysate was centrifuged at 12,500 rpm for 1 hour to remove insoluble materials. The supernatant was applied to anti-Flag M2 affinity resin (MilliporeSigma), washed with wash buffer (25 mM HEPES, pH 7.4, 150 mM NaCl, 0.02% (w/v) GDN (glyco-diosgenin, Anatrace)), and eluted using the wash buffer supplemented with 400 μg/mL Flag peptide. The eluent from the anti-Flag resin was loaded onto Strep-Tactin XT resin (IBA), washed with the wash buffer, and eluted using a buffer containing 25 mM HEPES (pH 7.4), 150 mM NaCl, 0.02% (w/v) GDN, and 50 mM D-biotin. The 3A2 complex was concentrated to 0.8 mL and further purified using size-exclusion chromatography (SEC) on a Superose 6 Increase 10/300 column (GE Healthcare). Western blot analysis was performed on SEC fractions using mouse monoclonal antibodies against the Flag tag (CW0287M, CWBIO) and Strep tag (ab76949, Abcam) for verification. The 3A3 complex was purified using similar procedures.

### Preparation of soluble Vγ9Vδ2 TCR

Plasmids encoding the soluble TCRγ9 and TCRδ2 chains (1 mg per plasmid) were pre-mixed with 4 mg of polyethyleneimine (PEI) in 45 mL of fresh medium for 20 minutes. This mixture was then transfected into ExpiHEK293F cells. The transfected cells were cultured at 37°C under 5% CO_2_ for 5 days before harvesting. Following centrifugation at 4000g for 20 minutes, the supernatant was collected and filtered through a 0.45 μm membrane to remove cell debris. The filtered medium was concentrated to approximately 150 mL and loaded onto Ni-NTA resin (Qiagen). The resin was extensively washed with a buffer containing 25 mM HEPES (pH 7.4), 150 mM NaCl, and 30 mM imidazole. The secreted Vγ9Vδ2 TCR was eluted using the wash buffer supplemented with 300 mM imidazole. The eluate was concentrated and further purified by SEC on a Superdex 200 Increase 10/300 GL column in a buffer containing 25 mM HEPES (pH 7.4) and 150 mM NaCl.

### Reconstitution of G115 and MOP TCR with 3A2 complex or 3A3 complex

The WT or mutant BTN complexes was incubated with an excess molar ratio of the Vγ9Vδ2 TCR to facilitate the formation of a stable complex. This incubation was performed at 4 °C for 1 hour with 40 µM HMBPP. The mixture was concentrated to a final volume of 0.8 mL. Any excess TCR was subsequently removed using SEC (Superose 6 increase 10/300, GE Healthcare) in a buffer containing 25 mM HEPES (pH 7.4), 150 mM NaCl, 0.02% (w/v) GDN, and 4 µM HMBPP. Finally, the purified TCR–BTN complex was concentrated and prepared for cryo-EM samples.

### Cryo-EM sample preparation and data collection

Holy carbon grids (Quantifoil, Au, 300-mesh, R1.2/1.3) were glow-discharged at 15 mA for 30 s. A 3 μL aliquot of concentrated sample was applied to the grid and blotted for 3 s under 100% humidity at 8°C. Subsequently, the grid was rapidly plunged into liquid ethane, using a Vitrobot Mark IV (Thermo Fisher Scientific).

For the cryo-sample of 3A2 complex with HMBPP, grids were transferred to a Krios electron microscopy (Thermo Fisher Scientific) operating at 300 kV and equipped with a Falcon 4i direct electron detector and a Selectris X energy filter. Movies in electron event representation (EER) format were automatically collected using EPU (Thermo Fisher Scientific) with a slit width of 10 eV and a defocus value range from −1.0 µm to −2.0 µm with a nominal magnification of 165,000X ^48^, corresponding a physical pixel size of 0.72 Å per pixel. Each movie was recorded for 4.98 s with a total dose of 50 e^−^ per Å^2^ contained in 1,530 EER frames.

For other cryo-samples, grids were transferred to a Krios electron microscopy (Thermo Fisher Scientific) operating at 300 kV and equipped with a Gatan K3 Summit detector and GIF Quantum energy filter. Movie stacks were automatically collected using EPU (Thermo Fisher Scientific) with a preset defocus value range from −1.0 µm to −2.0 µm in super-resolution mode with a nominal magnification of 81,000X^48^, corresponding to a physical pixel size of 1.087 Å per pixel or 1.0773 Å per pixel. Each movie was recorded for 2.56 s with a total dose of 50 e^−^ per Å^2^ contained in 32 frames.

### Cryo-EM sample preparation and data collection

For the dataset of 3A2 complex oligomer, the flowchart for data processing is presented in Figure S1E. Movie stacks were motion-corrected using MotionCor2 v1.2.6^49^, and the dose-weighted micrographs were subjected to Patch CTF Estimation in cryoSPARC v4^50^. The selection of micrographs was performed based on the aforementioned criteria. The template picking particles were extracted and binned by a factor of 4. The particles were subjected to multiple rounds of 2D classification to remove contaminations and junk particles. Cleaned particles were re-centered and re-extracted and binned by a factor of 2. After several rounds of 2D classification, the selected particles were subjected to Ab-Initio reconstruction (K=5). Particles from the best classes were combined and refined by non-uniform refinement^51^, yielding an oligomer map at 11.36 Å global resolution. The refined particles were not unsampled and further refined, as no discernible improvement in resolution or map quality was achieved. Validations and analysis of the density maps can be referred to Figures S1F and S1G.

The flowchart for data processing of 3A2 complex with HMBPP is presented in Figure S2C. EER movies were subjected to Patch Motion Correction and Patch CTF Estimation in cryoSPARC v4^50^. Micrographs exhibiting a CTF estimated resolution of worse than 4 Å, an astigmatism greater than 500, and a relative ice thickness greater than 1.1 were discarded by Manually Curate Exposure job. To perform template-based particle picking, a subset of motion-corrected micrographs was initially processed to generate 2D class averages with clear protein features. These class averages were subsequently utilized as references for template picking, which was conducted on the complete dataset. Extracted particles were binned by a factor of 4 initially and subjected to multiple rounds of 2D classification to discard junk particles. Cleaned particles were re-centered and re-extracted without downscaling and subjected to parallel Ab-Initio reconstruction (K=5). Particles from the best classes were merged and the duplicates were removed, and the resulting particles were refined using the best initial model as the reference. The refined particles were subsequently subjected to rounds of heterogeneous 3D classification, local motion correction, and non-uniform refinement ^51^, yielding a 2.96 Å overall reconstruction. The resulting particles were subjected to local refinement with a soft mask that encompassed the extracellular domain or the micelle plus intracellular domain, yielding local maps at 2.90 Å resolution and 2.73 Å resolution, respectively. Validations and analysis of the density maps can be referred to Figure S3.

For the dataset of 3A3 complex with HMBPP, the flowchart for data processing is presented in Figure S2D. Movie stacks were motion-corrected using MotionCor2 v1.2.6^49^, and dose-weighted micrographs were imported into cryoSPARC v4^50^. Following the aforementioned data processing strategies, the non-uniform refinement^51^ generated a 3.86 Å overall reconstruction. After that, those poses were transferred to RELION-5^52^ (https://github.com/3dem/relion/tree/ver5.0) using the csparc2star.py in the UCSF pyem program packages^53^ and then subjected to Bayesian polishing^54^. Bayesian polished particles were subjected to Relion 3D auto-refinement, yielding an overall density map with Fourier shell correlation (FSC) resolution of 3.81 Å, which exhibited superior map density compared to the map generated by cryoSPARC^50^. Subsequent consensus refinement with a soft mask that encompassed the extracellular domain (ECD) or the micelle plus intracellular domain (ICD), yielding two local maps at 3.47 Å resolution and 3.48 Å resolution, respectively. To improve the density of the B30.2 domain of BTN3A3, the resulting ICD refined particles were subjected to several rounds of focused 3D classification (skip alignment) with a sphere mask encompassing the targeted region. Subsequent consensus refinement on the sorted particles yielded a local density map at 3.26 Å resolution with superior B30.2 densities of BTN3A3 (Figure S2D). Validations and analysis of the density maps can be found in Figure S3. A similar data processing strategy was applied to the dataset of G115 TCR–3A2 complex (Figure S9D), MOP TCR–3A2 complex (Figure S9E), and G115 TCR–3A3 complex (Figure S9F), with the addition of HMBPP.

To resolve the density of the TCR only bound to the other side of the BTN2A1 IgV domain in G115 TCR–3A2 complex, we first generated a spherical mask centered on the interaction interface between the TCR and BTN2A1. This mask was subsequently applied in multiple rounds of 3D classification in RELION^52^. In each round, particles of good classes were merged and deduplicated from the last four iterations. The selected particles were then imported into cryoSPARC^50^ for homogeneous reconstruction followed by non-uniform refinement, yielding an overall density map at approximately 3.8 Å resolution. In this map, one TCR engaged both BTN2A1 and BTN3A2, while the other TCR bound only to the BTN2A1 IgV domain. To further improve the density of the TCR bound to the BTN2A1 IgV domain, the particles were re-centered to the BTN2A1-TCR interface and re-extracted. Subsequent local refinement with a protein mask produced a local density map with a resolution of 4.0 Å. To improve the resolution further, the corresponding particles were transferred back to RELION^52^ for multiple rounds of 3D classification (skip align) with a protein mask to remove poorly aligned particles and junk. The cleaning particles were then subjected to consensus refinement with a protein mask, which improved the resolution to 3.8 Å. A similar data processing strategy was applied to the dataset of G115 TCR–3A3 complex, with the addition of HMBPP.

For the dataset of G115 TCR–AAR complex, the flowchart for data processing is presented in Figure S14B. Movie stacks were motion-corrected using MotionCor2 v1.2.6^49^, and dose-weighted micrographs were imported into cryoSPARC v4^50^. Particles were picked using template picking and extracted. After multiple rounds of 2D classification, Ab-Initio reconstruction, non-uniform refinement, 3D classification and local refinement, the particles were subjected to particles subtraction to mask out the micelle and flexible intracellular domain. Next, the subtracted particles were subjected to local refinement with re-centering and a new fulcrum location specified, yielding a reconstruction at 3.25 Å resolution. Finally, this portion of particles were transferred to RELION-5^52^ (https://github.com/3dem/relion/tree/ver5.0) using the csparc2star.py in the UCSF pyem program packages^53^ and subjected to consensus refinement with a soft mask encompassed only the extracellular domain, yielded an ECD reconstruction at 3.20 Å resolution. Validations and analysis of the density maps can be referred to Figure S14.

The resolution of reconstruction was determined by the gold standard Fourier shell correlation using the FSC = 0.143 criterion in cryoSPARC v4^50^ or RELION-5^52^ (https://github.com/3dem/relion/tree/ver5.0). 3D Fourier shell correlation (FSC) analysis was conducted on the Remote 3DFSC Processing Server^55^ (https://3dfsc.salk.edu/). All figures depicting the maps were prepared using UCSF ChimeraX^56^ or UCSF Chimera^57^.

### Model building and refinement

The locally refined maps, sharpened by B-factors, were combined in UCSF Chimera^57^ to aid model building. The model of B30.2 domain of BTN2A1/3A1 in complex with HMBPP (PDB ID: 8JYE^34^), the model of BTN2A1 in complex with Vγ9Vδ2 TCR (PDB ID: 8DFW^58^), the AlphaFold2^59^ predicted models of BTN2A1 homodimer, BTN3A1/3A2 heterodimer and BTN3A1/3A3 heterodimer were used as initial models and rigidly fitted into corresponding density maps using USCF Chimera^57^, respectively. Several iterative rounds of real space refinement with secondary structure and geometry restraints in Phenix ^60^ and manual adjustment in COOT^61^ were performed until the model fit the density well and met the criteria. To clarify, we only built the atomic model of the TCR that engages both BTN2A1 and BTN3A2/A3, despite the observation of another TCR binding BTN2A1 exclusively.

To monitor overfitting, the resultant model was refined against one of the two independent half-maps (hereafter called “half map 1”) in Phenix^60^, and the refined model was then converted to a map utilizing EMAN2^62^. This map was subsequently used to compute the FSC with the “half map 1” and “half map 2”. These two FSC curves were referred to as “model_vs_half_map1” and “model_vs_half_map2”, respectively. Similarly, the resultant model was refined against the summed map of the two half-maps, and the refined model was used to compute the FSC curve with the summed map, which was referred to as “model_vs_summed map”. For each model, all three of these FSC curves were plotted. All models in this work were validated using the Phenix.molprobity^63^ tool (Table S1). UCSF Chimera^57^ and UCSF ChimeraX^56^ were used for structure visualization and figure preparation.

### Measurement of binding affinity

Surface plasmon resonance (SPR) was performed using a Biacore 1K (Cytiva) to assess the binding kinetics of WT, AAR, and AAA complexes with the G115 TCR. A CM5 chip was used to immobilize these complexes, achieving a surface density of approximately 5000 response units with 10 mM sodium acetate. The soluble Vγ9Vδ2 TCR complex was injected in a two-fold dilution series (27.43 μM to 0.85719 μM) at a flow rate of 30 μL/min, with a contact time of 180 seconds and a dissociation time of 300 seconds. The final response was calculated by subtracting signals from the control flow cell (BTNL3/L8) and blank injections. Data fitting was done using a 1:1 binding kinetics model with Biacore Insight Evaluation Software (version 5.0.18.22102, Cytiva), and analysis was performed using GraphPad Prism version 9.

### FLIM-FRET measurement and analysis

Full-length BTN2A1, BTN3A1, and BTN3A2 were fused and separated by P2A ribosome-skipping sites (GSGATNFSLLKQAGDVEENPGP) and subcloned into an optimized PCAG vector. Two strategies were employed to fuse fluorescent proteins. In the first strategy, CFP (moxCerulean3^64^) was fused to the amino terminus of BTN3A1, and YFP (mGold^65^) to BTN2A1. Constructs included: BTN2A1-*P2A*-CFP-BTN3A1-*P2A*-BTN3A2 (donor) and YFP-BTN2A1-*P2A*-CFP-BTN3A1-*P2A*-BTN3A2 (donor + acceptor). In the second strategy, CFP or YFP was fused to the carboxyl terminus of BTN3A2, generating BTN2A1-*P2A*-BTN3A1-*P2A*-BTN3A2-CFP (donor) and BTN2A1-*P2A*-BTN3A1-*P2A*-BTN3A2-YFP (acceptor).

Lenti-X 293T cells were transfected using Lipofectamine 3000 (Thermo Fisher Scientific). For the first strategy, donor plasmid was transfected alone (donor only group), and the donor + acceptor plasmid was transfected (donor + acceptor group). In the second strategy, donor plasmid was transfected alone (donor only group), while donor and acceptor plasmids were co-transfected at a 1:1 DNA mass ratio (donor + acceptor group). Cells were cultured for 14 hours before live-cell imaging.

Fluorescence lifetime measurements of the donor fluorophore were conducted with a Leica STELLARIS 8 microscope, featuring a temporal resolution of 97 ps, confocal scan head with FPGA electronics, 448 nm pulsed laser excitation, and fast spectral single-photon counting detectors. For FLIM-FRET analysis, FLIM images were generated by recording the average photon arrival per pixel. To ensure accuracy, a minimum of 2,000 photons per pixel were collected for both donor and acceptor acquisitions. The multi-exponential donor fit model was used to analyze time-correlated single-photon counting decay curves. The amplitude of each lifetime component was determined, and the intensity-weighted mean lifetime (τ) was calculated. For the donor-only group, CFP-expressing cells were analyzed, and the average weighted mean lifetime (τ) was measured using the n-Exponential Deconvolution model. In the donor + acceptor group, cells expressing both CFP and YFP were analyzed to determine the average lifetime within the cell membrane. FRET efficiency (E) was calculated using the equation:

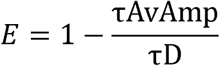

where τD is the lifetime of the unquenched donor, and τAvAmp is the amplitude-weighted average decay time.

### Analytical ultracentrifugation (AUC)

AUC analysis of purified protein samples was performed on Optima AUC-A/I analytical ultracentrifuge (Beckman Coulter). The complex was first purified by SEC and then concentrated to a final volume of 50 μL. Subsequently, samples were diluted to 380 μl in the AUC buffer (25 mM HEPES pH 7.4, 150 mM NaCl, 0.02% GDN) and loaded onto a double-sector sapphire cell, along with 400 μl reference buffer (25 mM HEPES pH 7.4, 150 mM NaCl). Sedimentation velocity analytical ultracentrifugation (SV-AUC) was carried out using an An-50 Ti rotor at 40,000 rpm and 20 °C. Sedimentation profiles were monitored by absorbance at 280 nm. Data were analyzed using SEDFIT. The extinction coefficient for the 3A2 complex was calculated using the Expasy server (https://www.expasy.org).

### Generation of NIH3T3 stable cell lines

For NIH3T3 stable cell lines expressing different BTN variants, BTN2A1, BTN3A1, and BTN3A2 were cloned into the lentiCRISPR v2 vector with a spleen focus-forming virus (SFFV) promoter. The HA tag and Flag tag were fused to N terminus of BTN2A1 and BTN3A1 separately, and along with a mcherry tag at the C terminus of BTN3A2, which were separated by a P2A ribosome-skipping site (ATNFSLLKQAGDVEENPGP). The lenti-plasmids along with the pMD2.G and psPAX2, were co-transfected into Lenti-X 293T cells to pack the lentivirus. Supernatants were collected 48 hours and 72 hours post-transfection. Then, the lentivirus was concentrated using 80 μg ml^−1^ protamine sulfate (Macklin) and 80 μg ml^−1^ chondroitin sulfate C (Macklin) before transduced into NIH3T3 cells. Transduced NIH3T3 cells exhibiting comparable mCherry fluorescence intensities were sorted via flow cytometry. Additionally, to ensure equivalent expression levels of the WT 3A2 complex, AAR mutant, and Y127A mutant, anti-Flag (BioLegend) or anti-HA tag antibodies (BioLegend) were employed.

### Multi-SIM imaging

To measure the oligomer formation and length distribution of 3A2 complex, NIH3T3 cells were transduced with pLVX-mGold-3A2 complex or indicated mutants with P_TRE3GS_ inducible promoter, which is regulated by doxycycline. NIH3T3 stable cell line was prepared using the same lentivirus packaging system. For super-resolution imaging of 3A2 complex and mutants, cells were seeded onto dishes (Cellvis) and were treated with doxycycline for 18h. The specimens were washed with PBS, then were fixed with 4% paraformaldehyde, followed by imaging in the TIRF-SIM mode. Images were visualized and processed by Fiji ImageJ. Statistical length distribution analysis of oligomer was performed using Prism (GraphPad).

### Vγ9Vδ2 T cell expansion

Cryopreserved human peripheral blood mononuclear cells were purchased from Shanghai Sailybio medical Co., LTD., which was approved by Shanghai Liquan Hospital Institutional Review Board (Project no. XF-WBC-220809). PBMCs (1 × 10^6^) were cultured in 12-well plates with 5 μM zoledronate treatment (Sigma). Fresh medium (OptiVitro® T cell medium, Excell Bio.) supplemented with human IL-2 and IL-15 (PeproTech) was replaced every 3 days. Vγ9Vδ2 T cells (purity > 90%) were expanded at 15 days and confirmed by anti-human CD3 (BD Pharmingen) and anti-human TCR Vδ2-FITC (Biolegend) antibodies. The protocol was approved by the Institutional Review Board of Westlake University (project no.20241010xwz).

### Functional assessment of human polyclonal Vγ9Vδ2 T cell activation assay

NIH3T3 cells expressing SFFV-3A2 complex or indicated mutants were seeded in 24-well plates for 12h, followed by co-incubation with in vitro-expanded Vγ9Vδ2 T cells from PBMCs (5 × 10^5^) ± zoledronate (5 μM). After 24 hours, culture supernatants were collected from the Vγ9Vδ2 T cells. The medium was measured for IFN<u>γ</u> expression levels using a human IFNγ ELISA kit (Beyotime), and absorbance was measured at 450 nm by a Multimode microplate reader (Varioskan LUX, Thermo Fisher Scientific). The Vγ9Vδ2 T cells were washed with FACS buffer and stained with anti-human TCR Vδ2-FITC and anti-human CD25-APC for 1h (BD Pharmingen). Flow cytometry analysis of CD25 expression was conducted on Cyto-FLEX LX-5L2 (Beckman). The raw data were analyzed using FlowJo (BD Biosciences).

### Tetramerization of soluble Vγ9Vδ2 TCR and Flow cytometry staining

The soluble Vγ9Vδ2 TCR with Avi tag was enzymatically biotinylated using biotin ligase BirA (made in house). The biotinylated Vγ9Vδ2 TCR was then incubated with Streptavidin-FITC (Solarbio) at a 4:1 molar ratio at 4 °C overnight. NIH3T3 cells expressing the WT 3A2 complex, Y127A mutant or AAR mutant were plated in 24-well plates and treated ± zoledronate for 24h. For Vγ9Vδ2 TCR tetramer staining, cells were digested by 0.25% Trypsin-EDTA, washed three times with FACS buffer, and subsequently incubated with 5 μg of Vγ9Vδ2 TCR-streptavidin complex at 4 °C for 1 hour. Data acquisition was performed using a Cyto-FLEX LX-5L2 flow cytometer (Beckman).

## Acknowledgments

This work is supported by National Key R&D Program of China (2020YFA0509300), Natural Science Foundation of Zhejiang Province (QKWL25C0501), National Natural Science Foundation of China (324B2025, 82241081), and Shenzhen Medical Academy of Research and Translation. We thank the cryo-EM facility, the high-performance computing center, the protein characterization and crystallography facility, and Y. Gao of the microscopy imaging platform of Westlake University for technical assistance, advice, and support. We thank Dr. Q. Huang for the generous gift of Lenti-X 293T.

## Author contributions

Q.Z., Q.S., and W.X. conceived the project and designed the experiments. W.X., Y.L. and Y.H. conducted cloning and protein purification. B.H. performed cryo-EM analysis. W.X., W.G. and W.Z. performed biochemical assays and other cell-based assays. All authors contributed to data analysis. Q.Z., Q.S., B.H. and W.X. wrote the manuscript. Q.Z. supervised the project.

## Data availability

All cryo-EM maps have been deposited in the Electron Microscopy Data Bank (EMDB; https://www.ebi.ac.uk/pdbe/emdb/), and all refined atomic models have been deposited in the Protein Data Bank (PDB; https://www.rcsb.org/). Accession numbers for all EMDB and PDB depositions are listed in the Table S1. All materials are available from the corresponding authors upon reasonable request.

## Supplementary figure titles and legends

**Figure S1.**
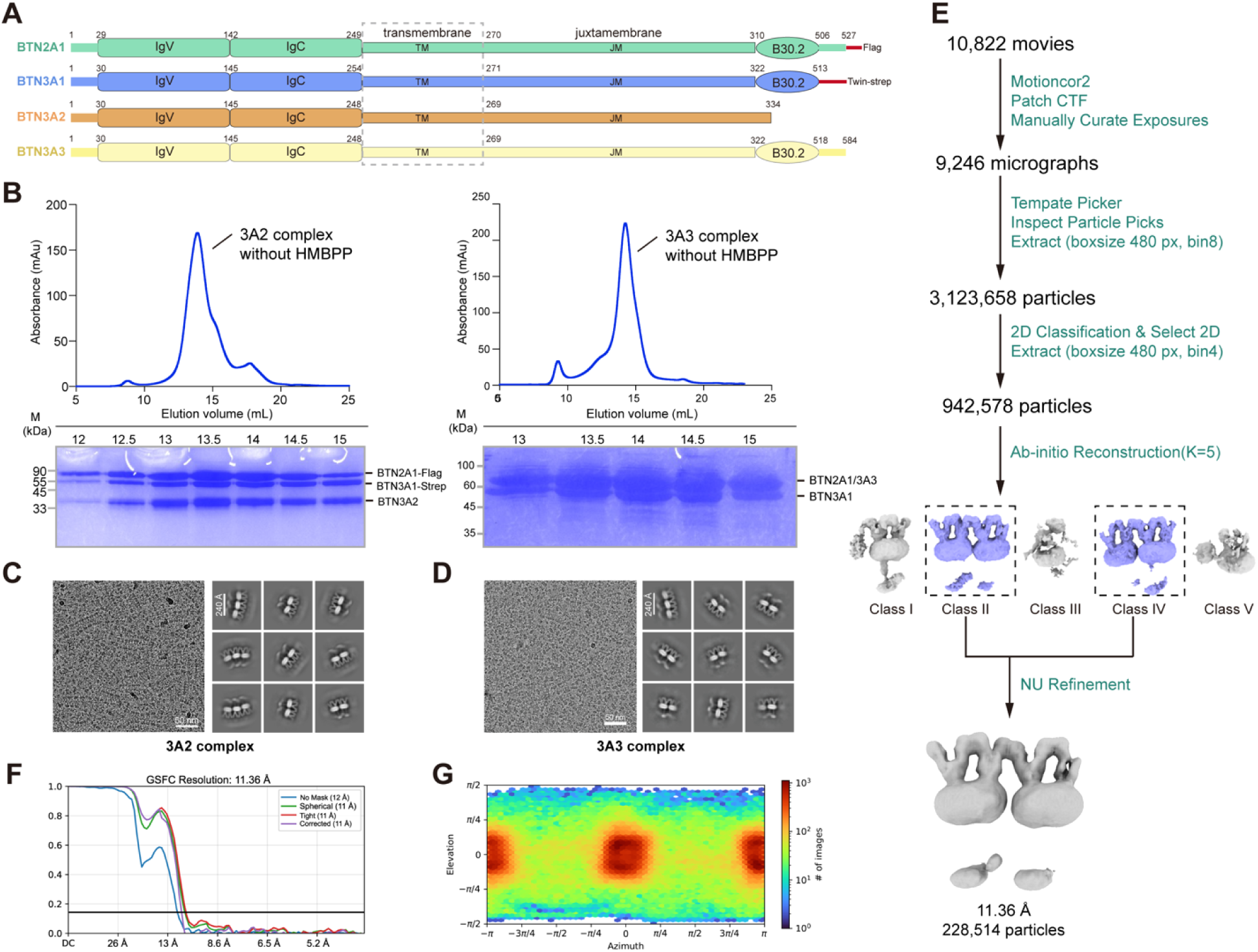
Biochemical characterization of BTN complex. (A) Schematic representation of domain organization of BTN2A1, BTN3A1, BTN3A2, and BTN3A3. TM: transmembrane; JM: juxtamembrane. Domain boundaries are indicated. (B) Representative size exclusion chromatography (SEC) profile of the 3A2 complex or 3A3 complex (left and right, top panel) and analysis of the relevant fractions by Coomassie blue staining (left and right, bottom panel) in the absence of HMBPP. (C and D) Representative cryo-EM image and 2D class averages of the 3A2 complex (C) or the 3A3 complex (D). (E) Summary of the cryo-EM data processing workflow for the 3A2 complex oligomer from (B). (F) FSC curves of the overall reconstruction of the 3A2 complex oligomer. (G) Angular distribution of particles used for the final reconstruction.

**Figure S2.**
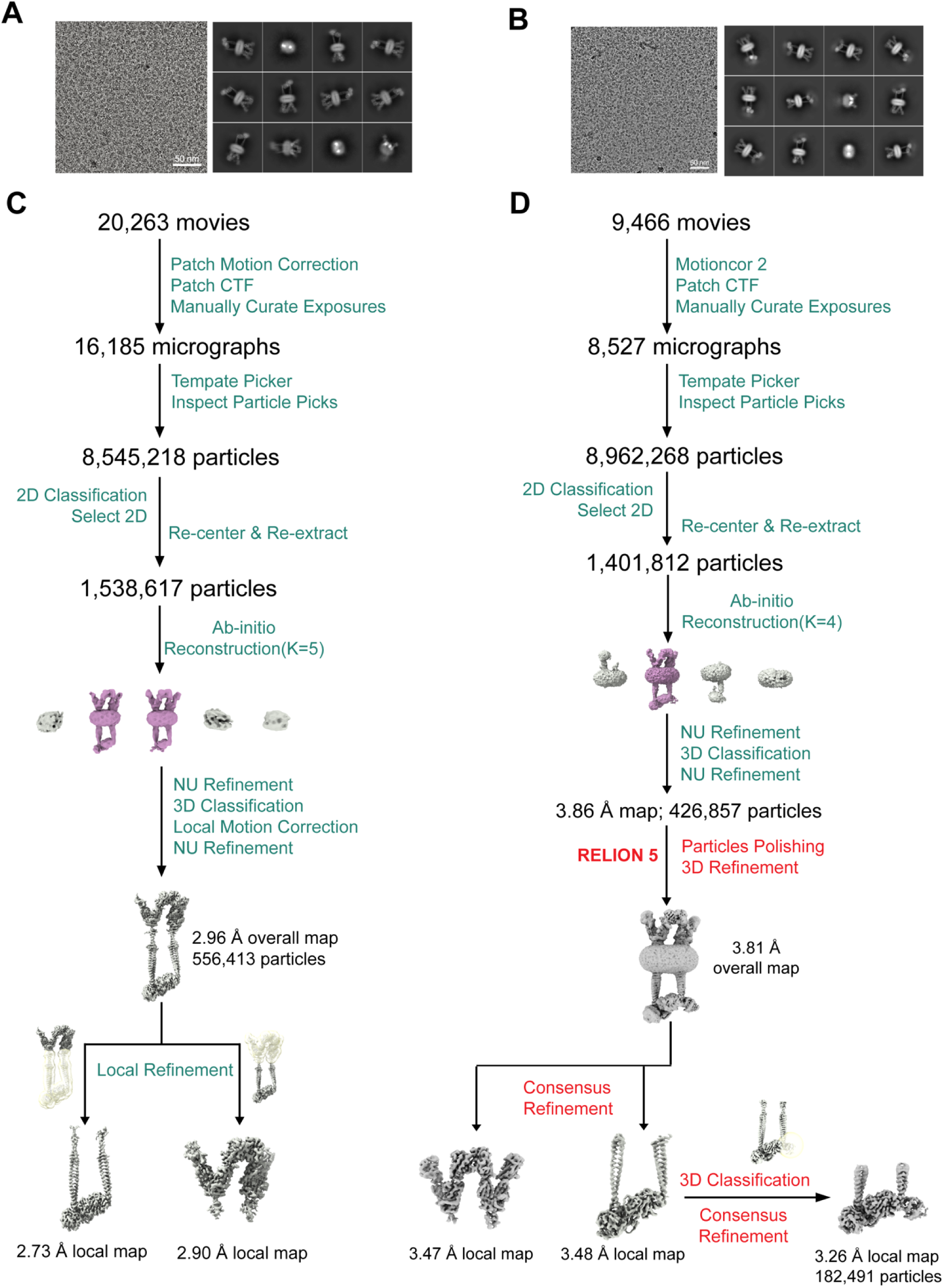
Cryo-EM analysis of pAg-bound BTN complexes. (A and B) Representative cryo-EM images and 2D class averages of the 3A2 complex (A) and the 3A3 complex (B) in the presence of 4 μM HMBPP. (C and D) Workflow of the cryo-EM data processing for the 3A2 complex (C) and the 3A3 complex (D) in the presence of HMBPP. Red annotations denote data processing steps conducted in RELION, whereas green annotations indicate computational procedures implemented using cryoSPARC.

**Figure S3.**
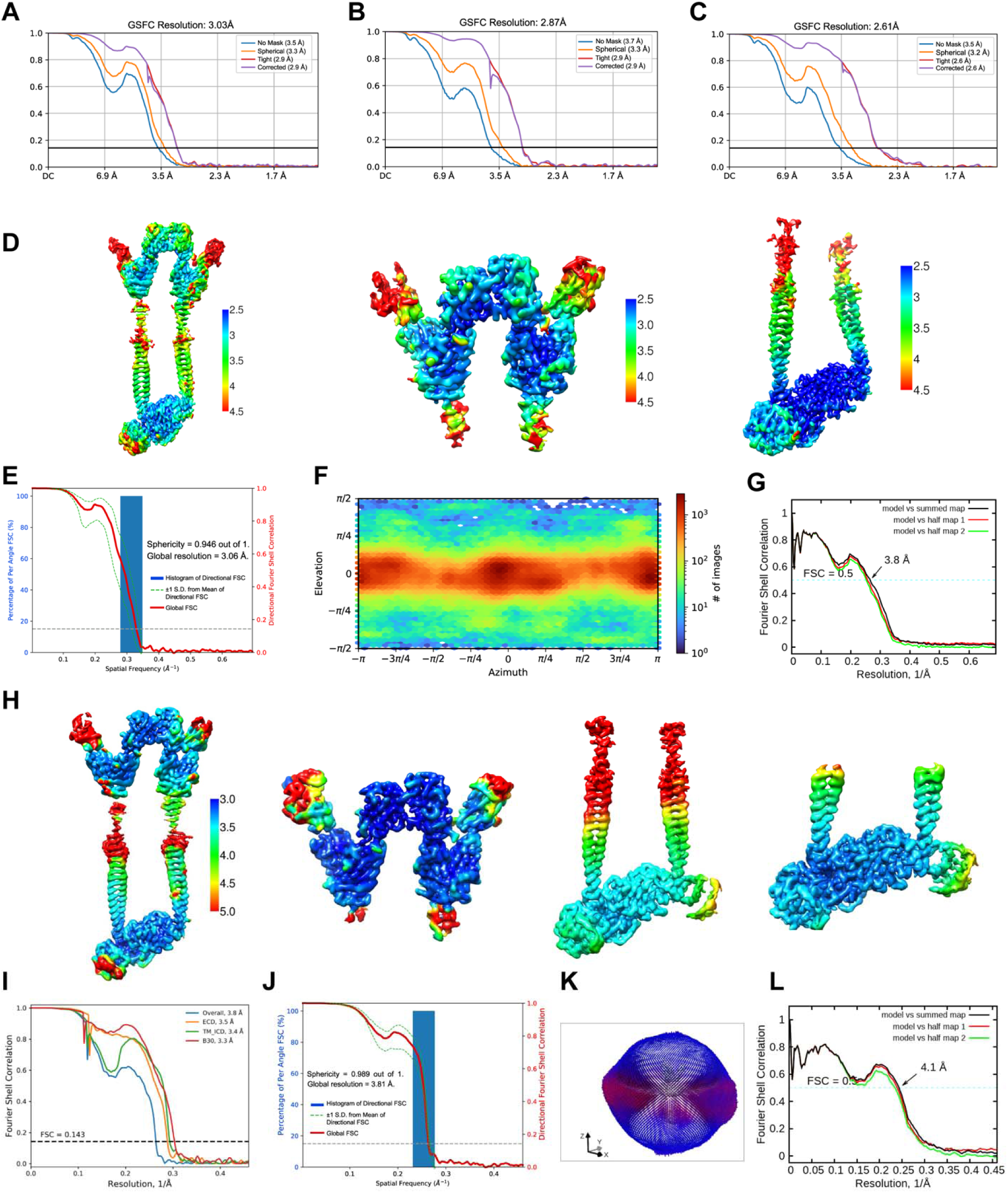
Validation of cryo-EM maps and atomic models of pAg-bound BTN complexes. (A-C) Fourier shell correlation (FSC) curves for the overall refinement (A), extracellular domain (ECD) local refinement (B), and intracellular domain (ICD) local refinement (C) of the 3A2 complex. (D) Local resolution estimation of the 3A2 complex overall map (left), ECD-focused map (middle), and ICD-focused map (right). Noise was manually removed for clarity. (E) The 3DFSC sphericity plot of the 3A2 complex overall map generated by 3DFSC. (F) Angular distribution of particles for the final reconstruction of the overall 3A2 complex. (G) Model versus map FSC curves for the 3A2 complex. (H) Local resolution estimation of the 3A3 complex overall map (left), ECD-focused map (middle), ICD-focused map (right), and B30.2-focused map (far right). Noise was manually removed for clarity. (I) FSC curves for overall refinement and consensus refinements of the 3A3 complex. Dashed line indicates FSC = 0.143. (J) The 3DFSC sphericity plot of 3A3 complex overall map generated by 3DFSC. (K) Angular distribution of particles for the final reconstruction of the overall 3A3 complex. (L) Model versus map FSC curves for the 3A3 complex. See Methods for specific details regarding FSC curve computation in (G) and (I).

**Figure S4.**
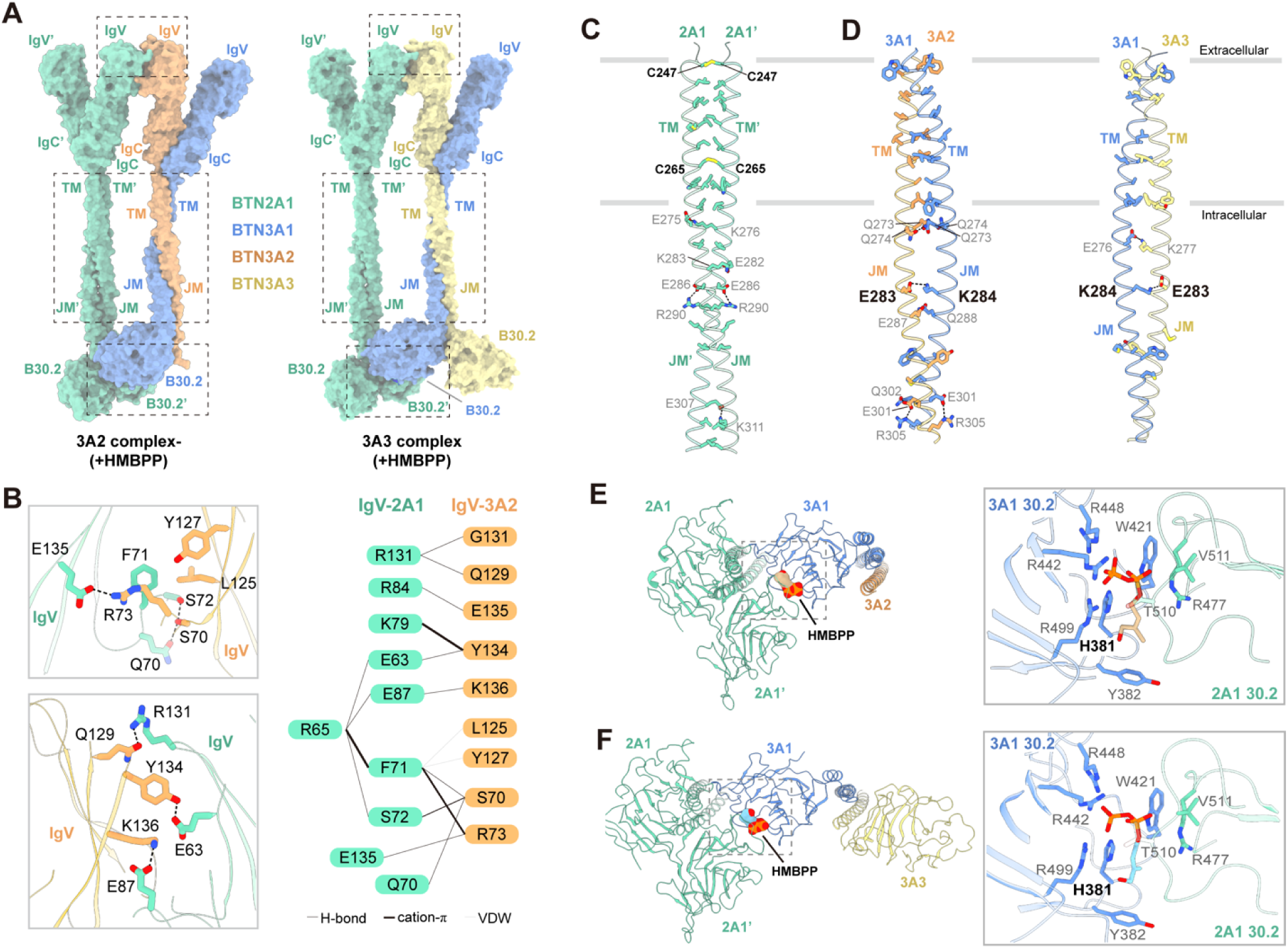
Interchain interactions determine BTN complex assembly. (A) Surface representation of the pAg-bound 3A2 complex (left) or the pAg-bound 3A3 complex (right). Dashed boxes highlight critical regions for BTN complex assembly. Subunit rendering: BTN2A1 (medium aquamarine), BTN3A1 (cornflower blue), BTN3A2 (sandy brown), and BTN3A3 (khaki). This color scheme is consistent throughout the work. (B) Close-up views of the IgV domain interface between BTN2A1 and BTN3A2 (left). Residues are shown as sticks, with hydrogen bonds, salt bridges, and Cation-π interactions indicated by dashed lines. The intricate interaction network between IgV_BTN2A1_ and IgV_BTN3A2_ is shown through a connection diagram of related residues (right). (C and D) Close-up views of the interface interactions between the TM/JM domains of the BTN2A1 homodimer (C), BTN3A1/3A2 heterodimer (D, left), and BTN3A1/3A3 heterodimer (D, right). Models are shown in cartoon style, with interacting residues depicted as sticks. Hydrogen bonds are dashed, and disulfide bonds are highlighted in yellow. TM: transmembrane; JM: juxtamembrane. (E and F) HMBPP binds to the B30.2 domain of BTN2A1 and BTN3A1 B30.2 domains within the 3A2 complex (E, left) or the 3A3 complex (F, left). The HMBPP is shown in sphere style. Close-up views of the HMBPP binding pocket within the BTN2A1 and BTN3A1 domains (E and F, right).

**Figure S5.**
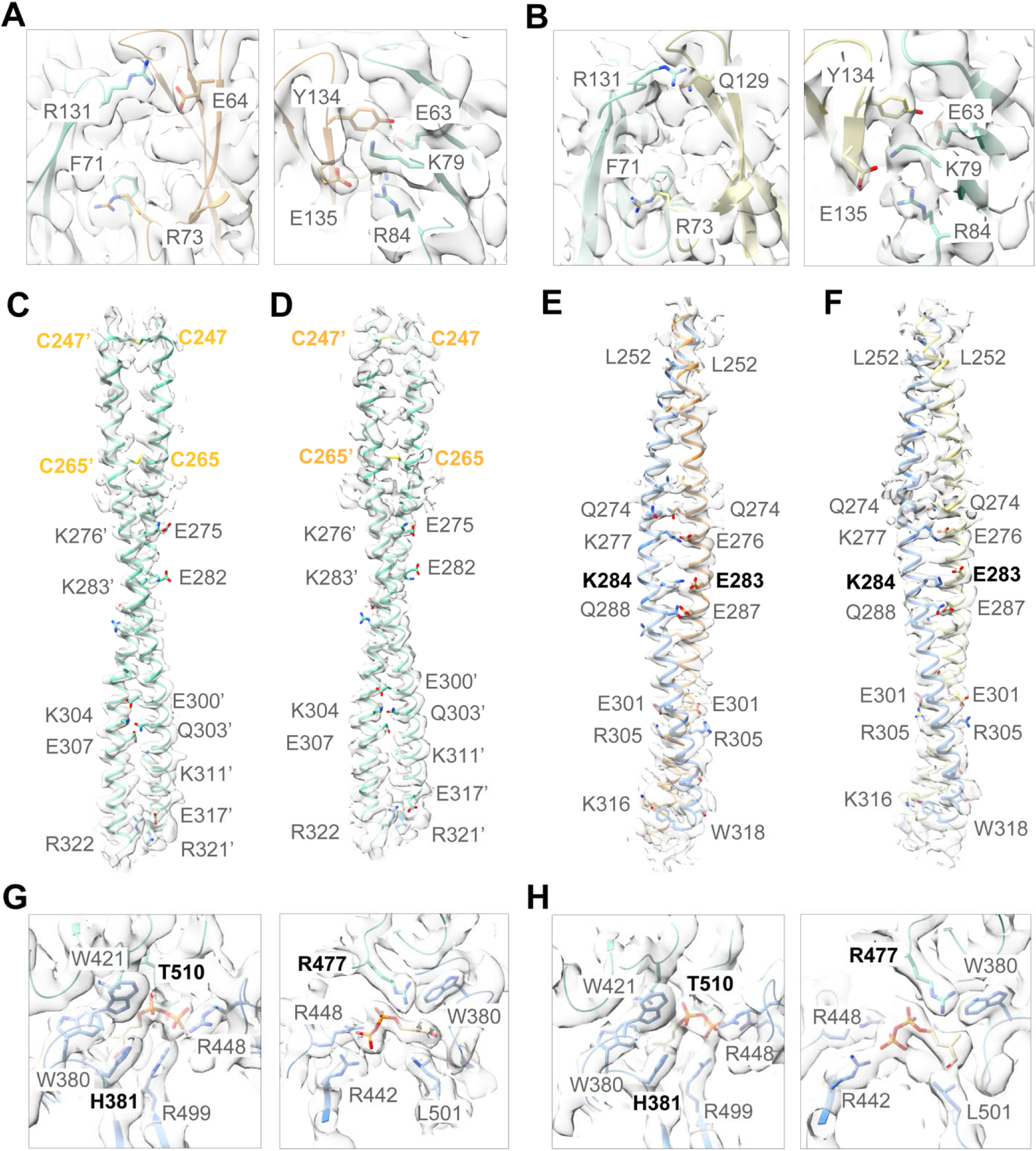
Representative cryo-EM maps of pAg-bound BTN complexes. (A) Close-up view of the interface between IgV_BTN2A1_ and IgV_BTN3A2_, shown in two views, with the map contoured at 8σ. (B) Close-up view of the interface between IgV_BTN2A1_ and IgV_BTN3A3_, shown in two views, with the map contoured at 8σ. (C and D) Close-up views of the interface between the TM/JM_BTN2A1_ and TM/JM_BTN2A1’_ regions in the 3A2 complex (C) and 3A3 complex (D), with the maps contoured at 6σ. (E and F) Close-up views of the interface between the TM/JM_BTN3A1_ and TM/JM_BTN3A2_ or TM/JM_BTN3A3_ regions in the 3A2 complex (E) and 3A3 complex (F), with the maps contoured at 8σ. (G and H) Close-up views of the HMBPP binding pockets in the 3A2 complex (G) and 3A3 complex (H), shown in two views, with the maps contoured at 11σ. Side chains of residues involved in bridging interactions are labeled and shown as sticks.

**Figure S6.**
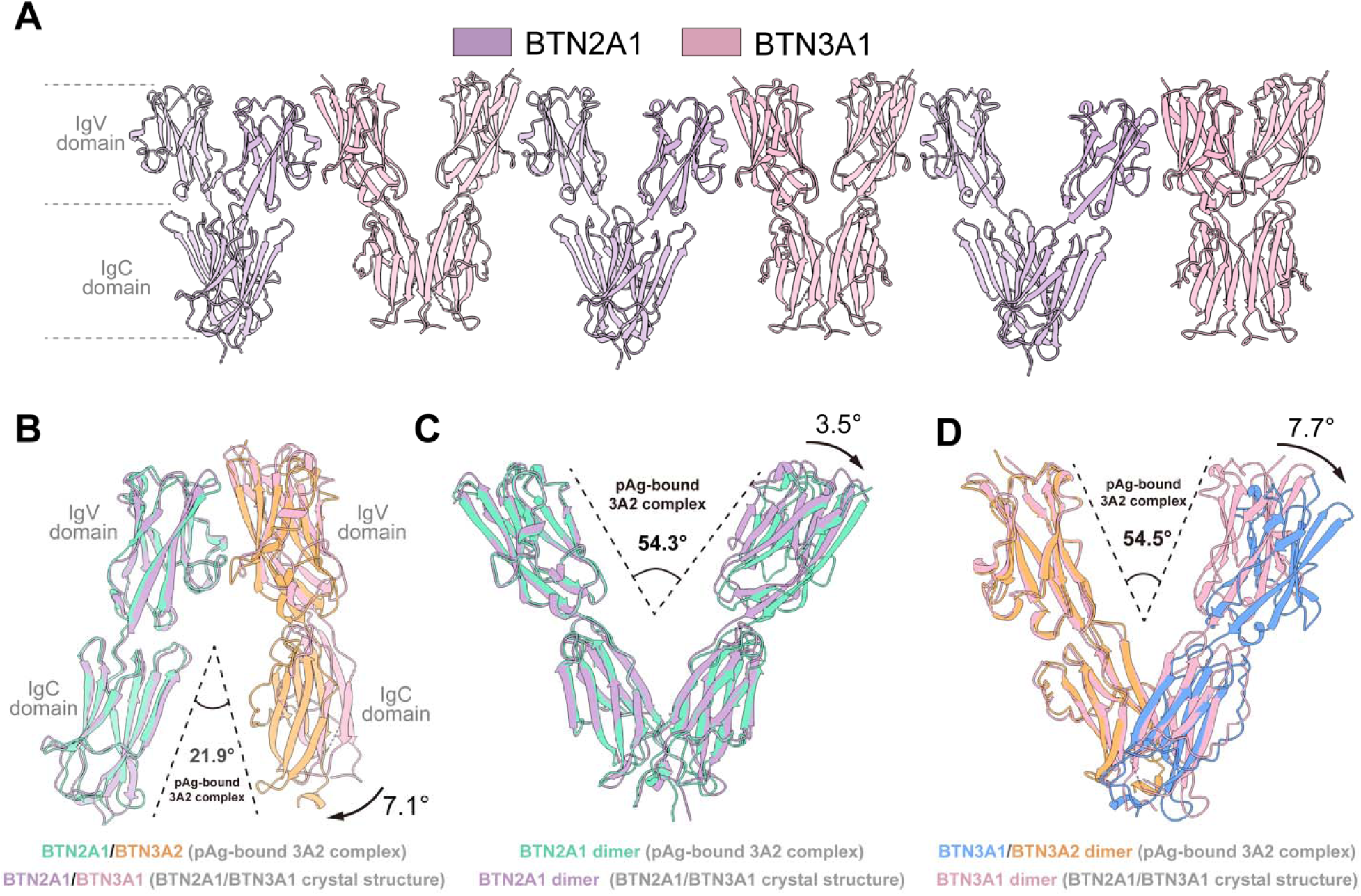
Structural comparison of the extracellular regions between the pAg-bound 3A2 complex and the BTN2A1/BTN3A1 complex. (A) The high-order oligomer (PDB ID: 8DFX^33^) in the crystal structure of BTN2A1 and BTN3A1 extracellular domain complex. (B) Comparison of the interchain angle between the Ig-like domains of BTN2A1 and BTN3A1. In the pAg-bound 3A2 complex, the BTN2A1/BTN3A1 angle is decreased by 7.1° compared to that in the crystal structure of BTN2A1 and BTN3A1 extracellular domain complex. (C) Comparison of the dimerization angle in the extracellular region of the BTN2A1 dimer. The BTN2A1 dimer angle in the pAg-bound 3A2 complex is increased by 3.5° relative to that of the BTN2A1 dimer in the crystal structure of BTN2A1 and BTN3A1 extracellular domain complex. (D) Comparison of the dimerization angle in the extracellular region of the BTN3As dimer. The BTN3A1/BTN3A2 angle in the pAg-bound 3A2 complex is increased by 7.7° compared to that of the BTN3A1 dimer in the crystal structure of BTN2A1 and BTN3A1 extracellular domain complex.

**Figure S7.**
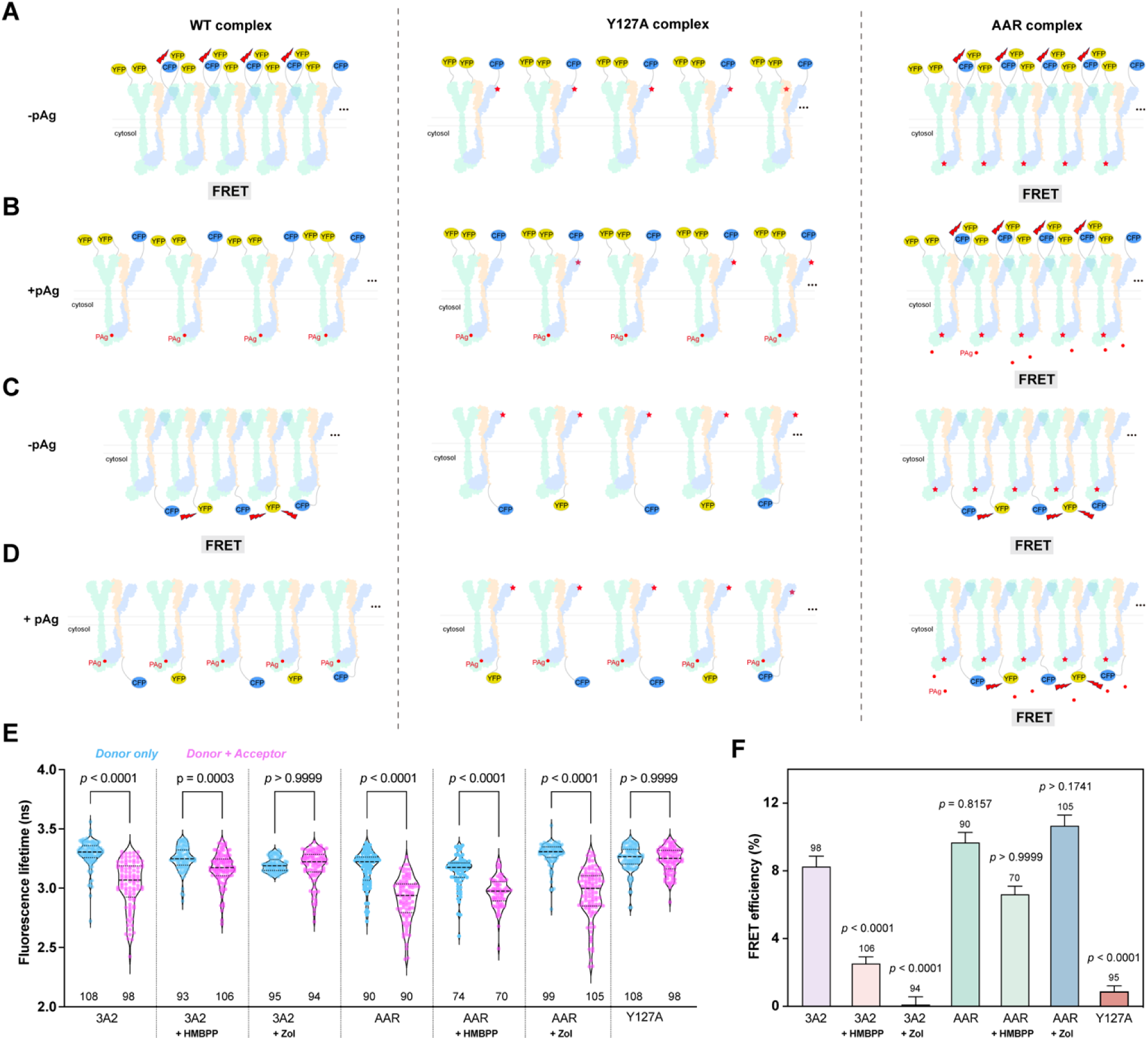
Design of FLIM-FRET assays to evaluate the oligomerization state of the 3A2 complex on cell membrane. (A and B) CFP and YFP are fused to the N termini of BTN3A1 and BTN2A1, respectively. In the absence of pAg, FRET occurs in both the WT and AAR complexes (A), while FRET is diminished in the WT complex but still occurs in the AAR complex in the presence of pAg (B). (C and D) CFP and YFP are fused to the C termini of BTN3A2. FRET occurs in both the WT and AAR complexes in the absence of pAg (C). In contrast, FRET is diminished in the WT complex but still occurs in the AAR complex (D). The mutations are indicated by red pentagrams. (E) Fluorescence lifetime of C-terminal CFP was measured in Lenti-X 293T cells using FLIM-FRET. “Donor only” refers to cells transfected with C-terminal CFP-labeled BTN3A2. “Donor + Acceptor” refers to cells co-expressing both C-terminal CFP-labeled and C-terminal YFP-labeled BTN3A2. The N value is provided for each group (mean ± SEM).(F) FRET efficiency was determined in cells expressing WT 3A2 complex and its variant, using the same constructs as in (E). The mean fluorescence lifetime in each “Donor only” group was used to calculate FRET efficiency in each “Donor + Acceptor” group. The N value is provided for each group (mean ± SEM). *P* values were calculated using the Kruskal-Wallis test with Dunn’s multiple comparisons post-test in (E and F). Y127A, the 3A2 complex variant harboring a BTN3A1-Y127A mutation to disrupt the oligomeric assembly; AAR, the 3A2 complex variant harboring BTN3A1-H381R/BTN2A1-R477A/T510A mutations to abolish HMBPP binding. Zol, zoledronate.

**Figure S8.**
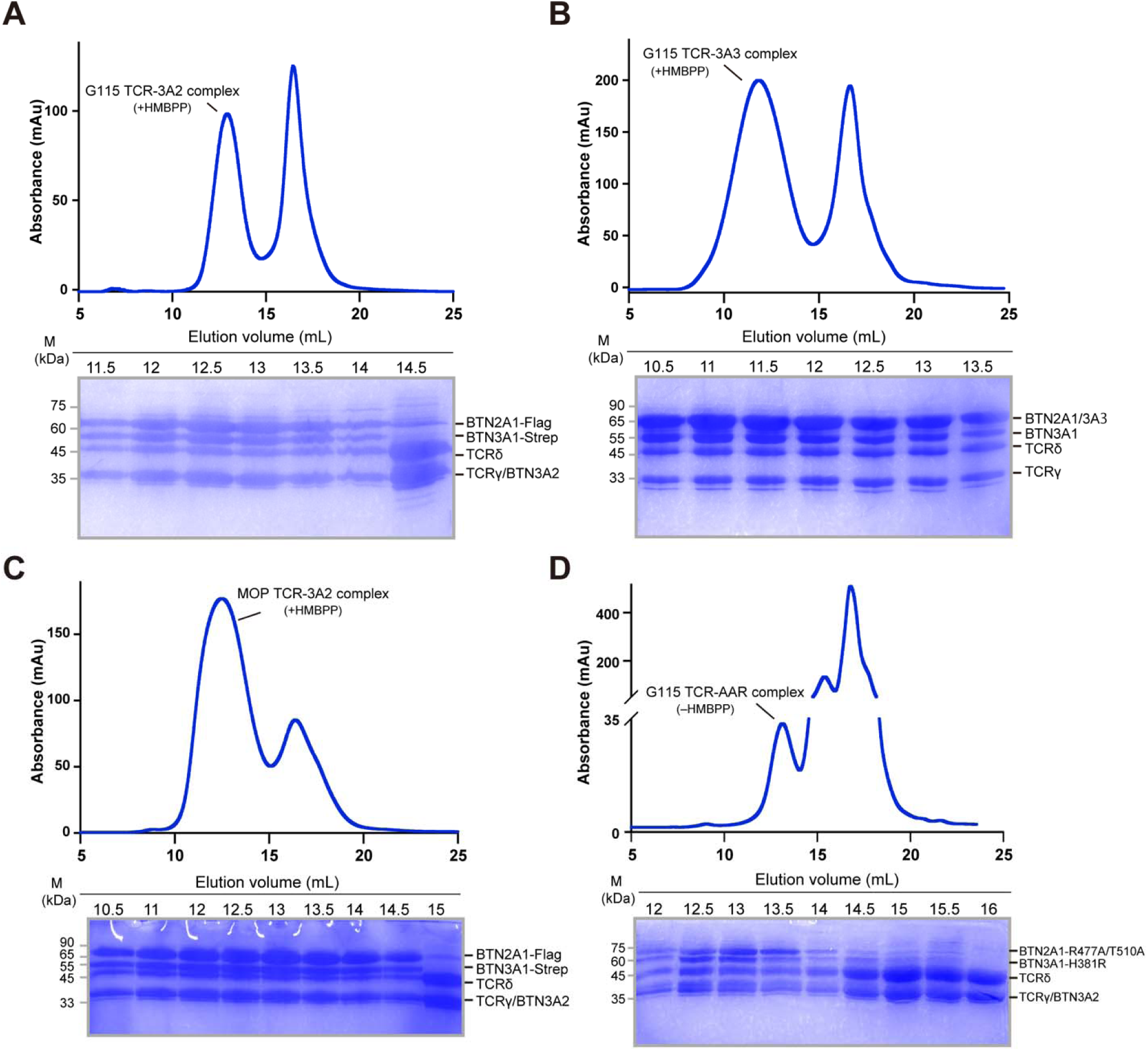
Biochemical characterization of the pAg-bound TCR–BTN complex and the TCR–AAR complex. (A–D) Representative size exclusion chromatography (SEC) profiles (top panels) and Coomassie blue staining analyses of corresponding fractions (bottom panels) for the pAg-bound G115 TCR– 3A2 complex (A), the pAg-bound G115 TCR–3A3 complex (B), the pAg-bound MOP TCR– 3A2 complex (C), and the G115 TCR–AAR complex (D). AAR denotes the 3A2 complex variant containing BTN3A1-H381R, BTN2A1-R477A, and T510A mutations to eliminate HMBPP binding. HMBPP was added at a final concentration of 4 μM in (A–C). See Methods for details on protein purification.

**Figure S9.**
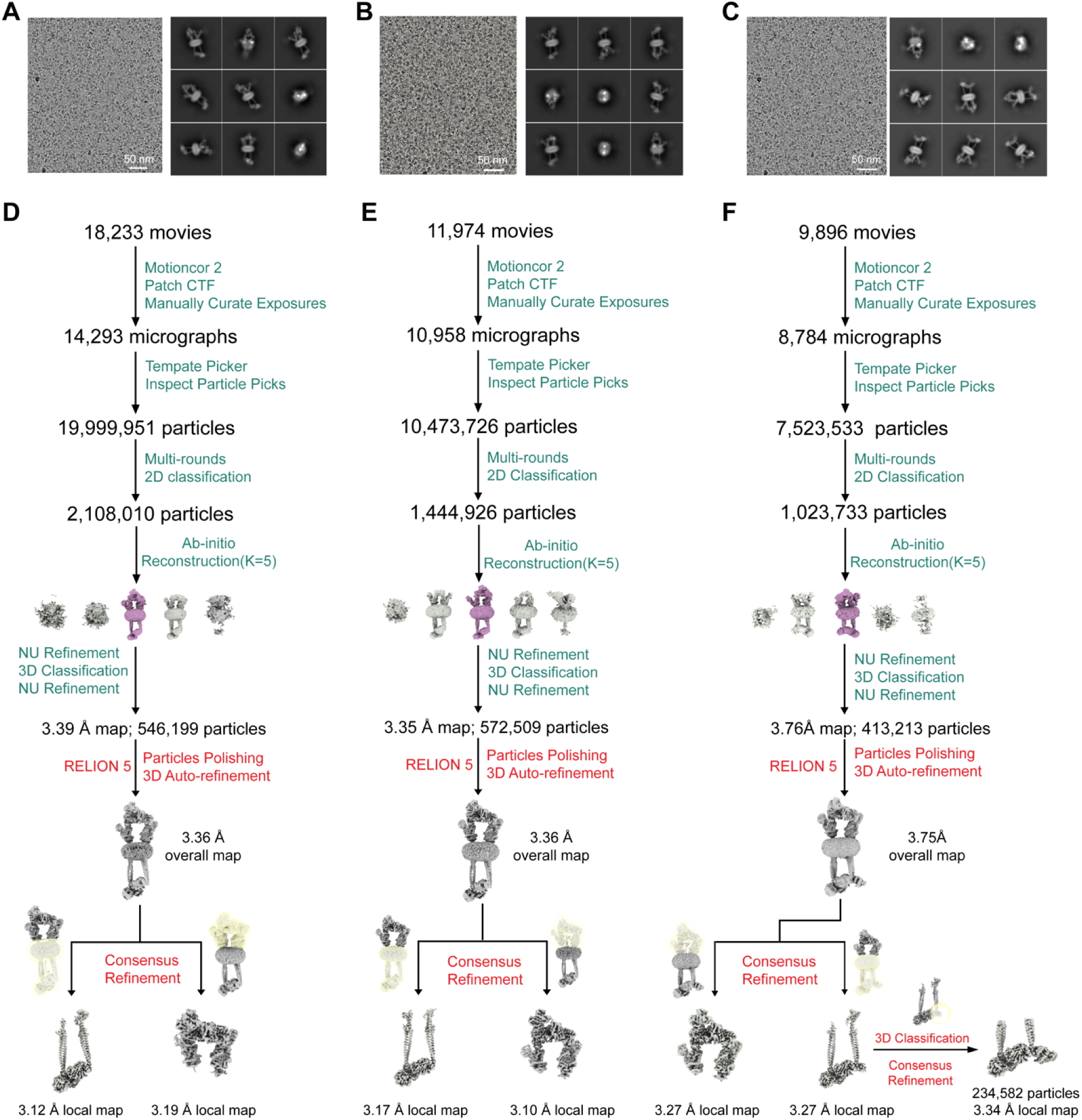
Cryo-EM analysis of pAg-bound Vγ9Vδ2 TCR–BTN complexes. (A–C) Representative cryo-EM images and 2D class averages of the pAg-bound G115 TCR– 3A2 complex (A), pAg-bound MOP TCR–3A2 complex (B), and pAg-bound G115 TCR–3A3 complex (C). (D–F) Workflow of the cryo-EM data processing for the pAg-bound G115 TCR–3A2 complex (D), the pAg-bound MOP TCR–3A2 complex (E), and the pAg-bound G115 TCR–3A3 complex (F).

**Figure S10.**
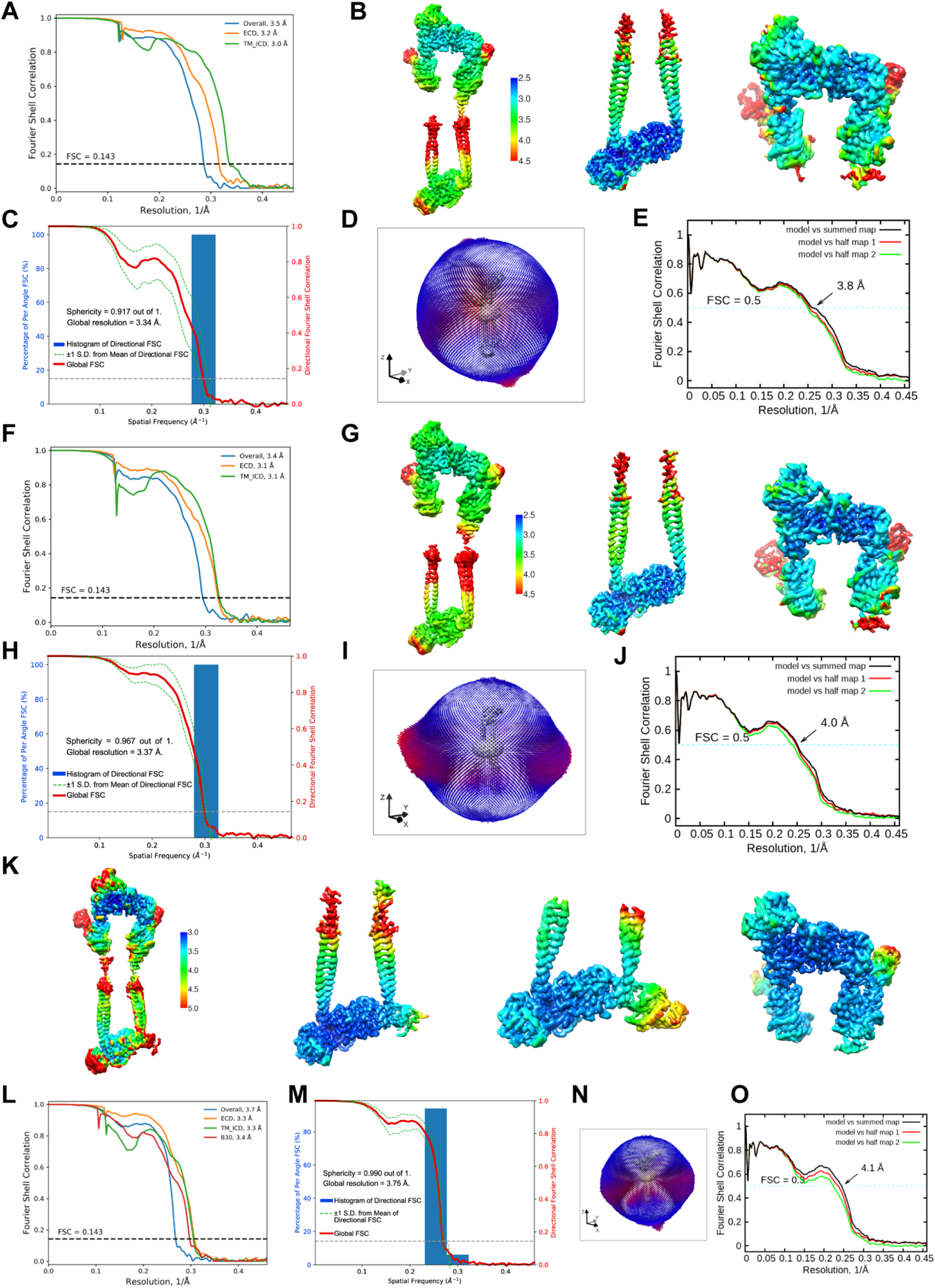
Validation of cryo-EM maps and models of pAg-bound Vγ9Vδ2 TCR–BTN complexes. (A) FSC curves of overall refinement and consensus refinements for the pAg-bound G115 TCR– 3A2 complex. (B) Local resolution estimation for the pAg-bound G115 TCR–3A2 complex overall map (left), ICD-focused map (middle), and ECD-focused map (right). Noise was manually removed for clarity. (C) 3DFSC sphericity plot generated by 3DFSC for the overall pAg-bound G115 TCR–3A2 complex. (D) Angular distribution of particles for the final reconstruction. (E) Model versus map FSC curves of the overall pAg-bound G115 TCR–3A2 complex. (F) FSC curves of overall refinement and consensus refinements for the pAg-bound MOP TCR– 3A2 complex. (G) Local resolution estimation for the MOP TCR–3A2 complex overall map (left), ICD-focused map (middle), and ECD-focused map (right). Noise was manually removed for clarity. (H) 3DFSC sphericity plot generated by 3DFSC for the overall pAg-bound MOP TCR–3A2 complex. (I) Angular distribution of particles for the final reconstruction. (J) Model versus map FSC curves for the MOP TCR–3A2 complex. (K) FSC curves of overall refinement and consensus refinements for the pAg-bound G115 TCR– 3A3 complex. (L) Local resolution estimation for the G115 TCR–3A3 complex overall map (left), ICD-focused map (middle), and ECD-focused map (right). Noise was manually removed for clarity. (M) 3DFSC sphericity plot generated by 3DFSC for the overall pAg-bound G115 TCR–3A3 complex. (N) Angular distribution of particles for the final reconstruction. (O) Model versus map FSC curves for the MOP TCR–3A3 complex.

**Figure S11.**
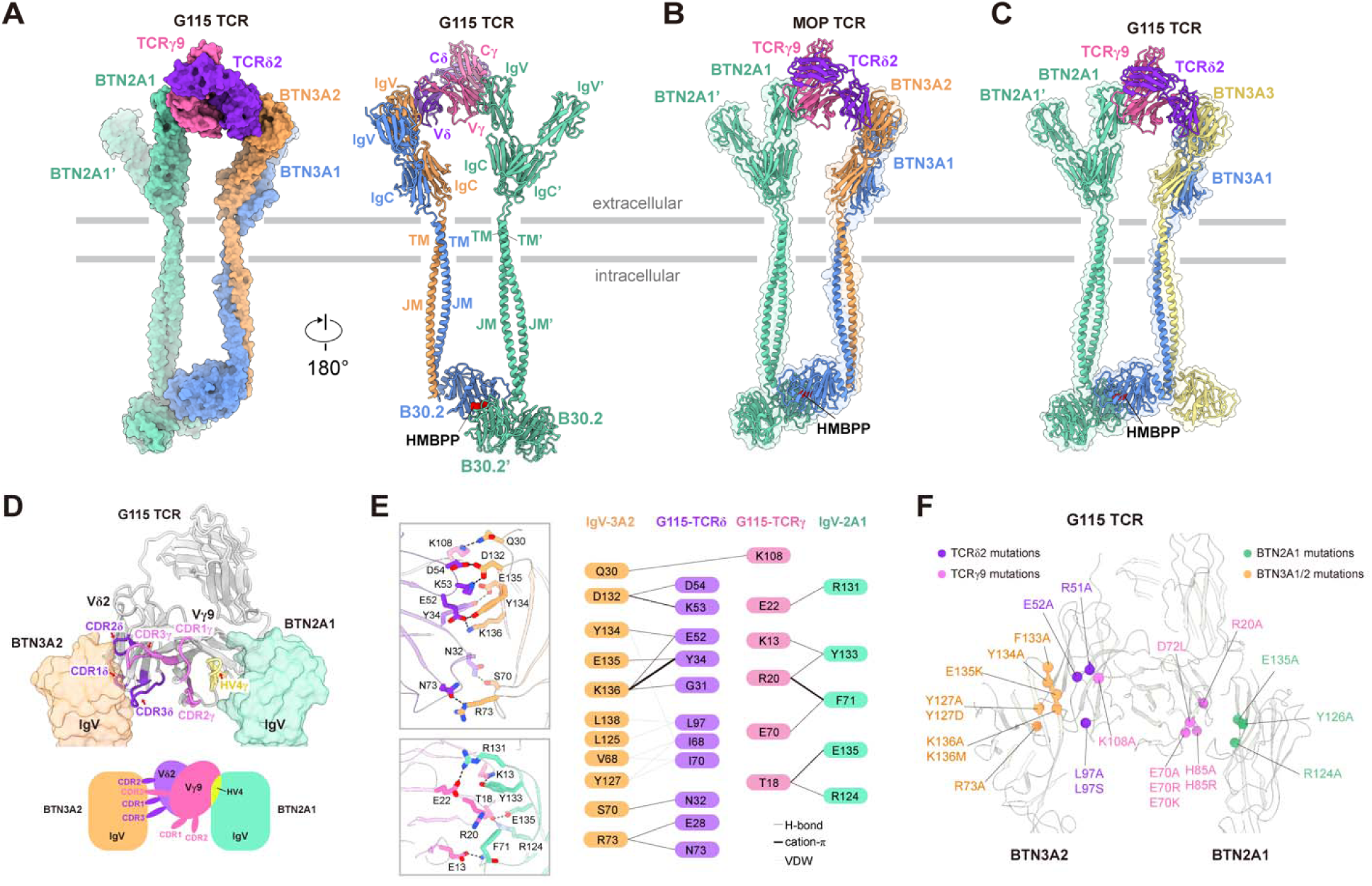
Vγ9Vδ2 TCR binds to the ectodomain of the BTN complex involving CDR loops and HV4 region. (A) Overall structure of Vγ9Vδ2 TCR (G115 prototype) in complex with the pAg-bound 3A2 complex (Figure 3A, right). The variable domains of the TCR are intricately positioned at the IgV_BTN2A1_–IgV_BTN3A2_ interface. The structure is shown as a surface (left) or cartoon (right), with HMBPP depicted as a red sphere. (B) Overall structure of Vγ9Vδ2 TCR (MOP prototype) in complex with the pAg-bound 3A2 complex. BTN molecules are shown as cartoons in the transparent surface, with TCR presented as cartoons and HMBPP shown as red sphere. (C) Overall structure of Vγ9Vδ2 TCR (G115 prototype) in complex with the pAg-bound 3A3 complex. Subunit colors are as follows: BTN2A1 (medium aquamarine), BTN3A1 (cornflower blue), BTN3A2 (sandy brown), BTN3A3 (khaki), TCRδ (blue violet), TCRγ (hot pink). (D) The CDR1, CDR2, CDR3, and HV4 regions of the G115 TCR are involved in recognizing the 3A2 complex. The CDRδ, CDRγ, and HV4γ regions are colored blue violet, hot pink, and yellow, respectively. (E) Close-up views of the interfaces between the 3A2 complex and G115 TCR (left two panels). The intricate interaction network is depicted by a connection diagram of related residues (right panel). (F) Mapping of mutations onto the G115 TCR–3A2 complex structure.

**Figure S12.**
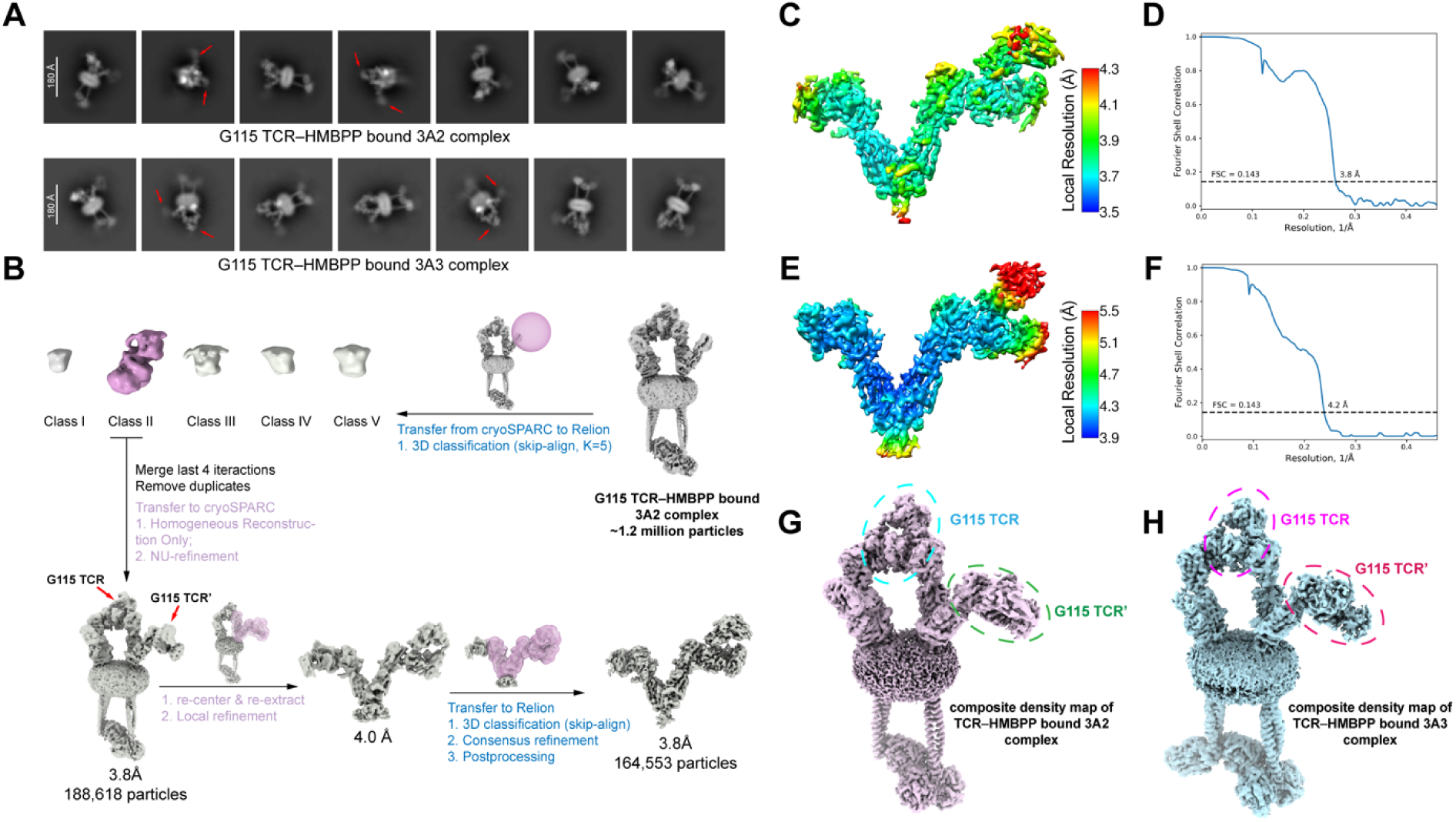
Cryo-EM data processing and analysis of the two Vγ9Vδ2 TCRs engaging the ectodomain of the pAg-bound BTN complexes. (A) Selective 2D class averages indicate two TCRs bind ectodomains. Upper panel: the G115 TCR–HMBPP bound 3A2 complex; lower panel: the G115 TCR–HMBPP bound 3A3 complex. (B) Data processing workflows for the G115 TCR–HMBPP bound 3A2 complex. A similar workflow was applied to the dataset of the G115 TCR–HMBPP bound 3A3 complex. (C and D) Local resolution estimation (C) and Fourier shell correlation (FSC) curves (D) for the G115 TCR–HMBPP bound 3A2 complex. The map is contoured at 9σ. (E and F) Local resolution estimation (E) and Fourier shell correlation (FSC) curves (F) for the G115 TCR–HMBPP bound 3A3 complex. The map is contoured at 7σ. (G and H) Composite density maps of the G115 TCR–HMBPP bound 3A2 complex contoured at about 7σ (G) and G115 TCR–HMBPP bound 3A3 complex contoured at about 7σ (H). The additional locally refined maps used for combination are shown in Figure S9.

**Figure S13.**
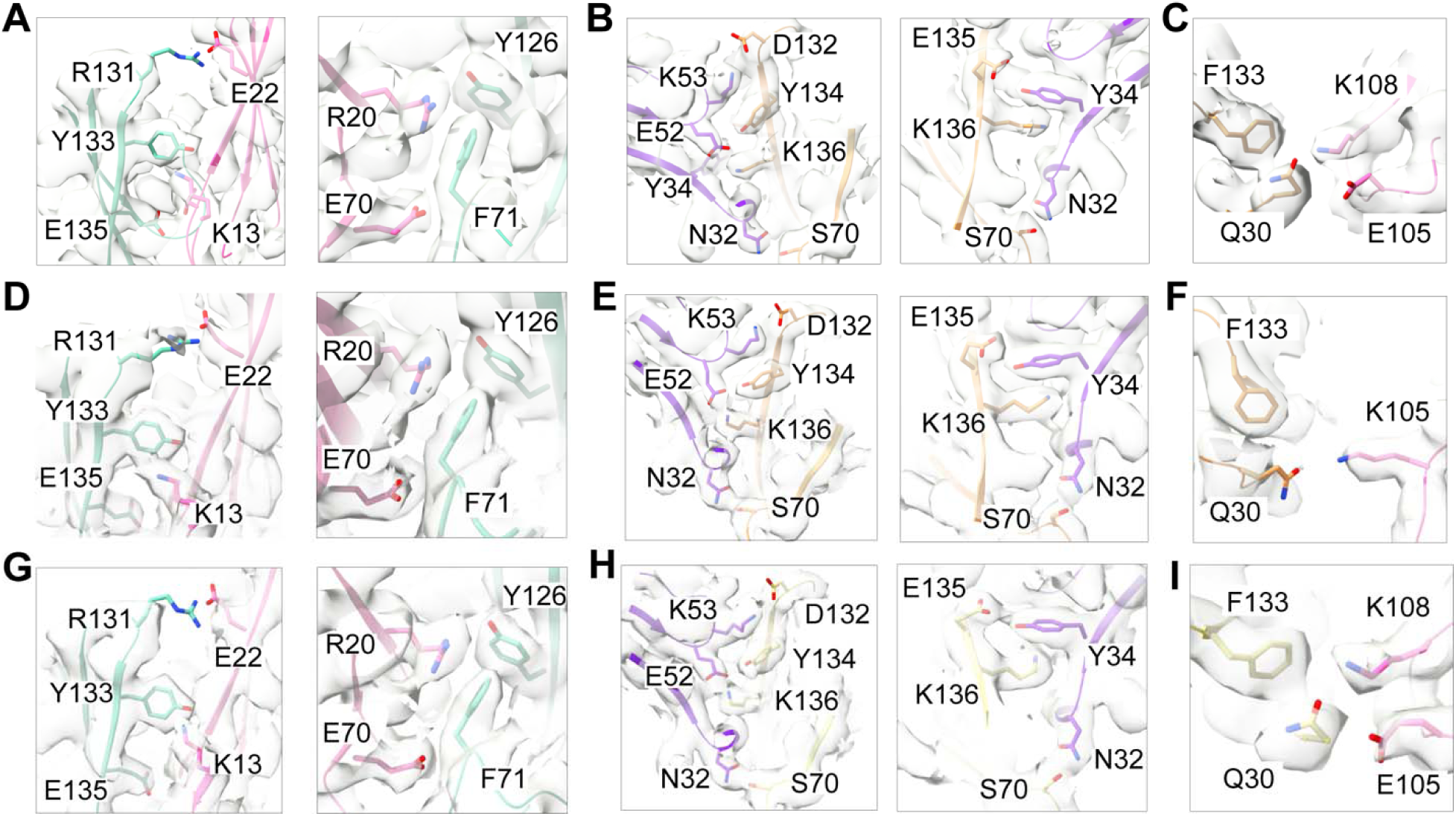
Representative cryo-EM maps of pAg-bound Vγ9Vδ2 TCR–BTN complexes. (A–C) The interface between G115 TCR and the 3A2 complex: Close-up views of the interface between IgV_BTN2A1_ and the Vγ9 domain (A), IgV_BTN3A2_ and the Vδ2 domain (B), and IgV_BTN3A2_ and the Vγ9 domain (C). (D–F) The interface between MOP TCR and the 3A2 complex: Close-up views of the interface between IgV_BTN2A1_ and the Vγ9 domain (D), IgV_BTN3A2_ and the Vδ2 domain (E), and IgV_BTN3A2_ and the Vγ9 domain (F). (G–I) The interface between G115 TCR and the 3A3 complex: Close-up views of the interface between IgV_BTN2A1_ and the Vγ9 domain (G), IgV_BTN3A3_ and the Vδ2 domain (H), and IgV_BTN3A3_ and the Vγ9 domain (I). The map is contoured at 9σ (A–C), 6σ (D–F), and 9σ (G–I), respectively.

**Figure S14.**
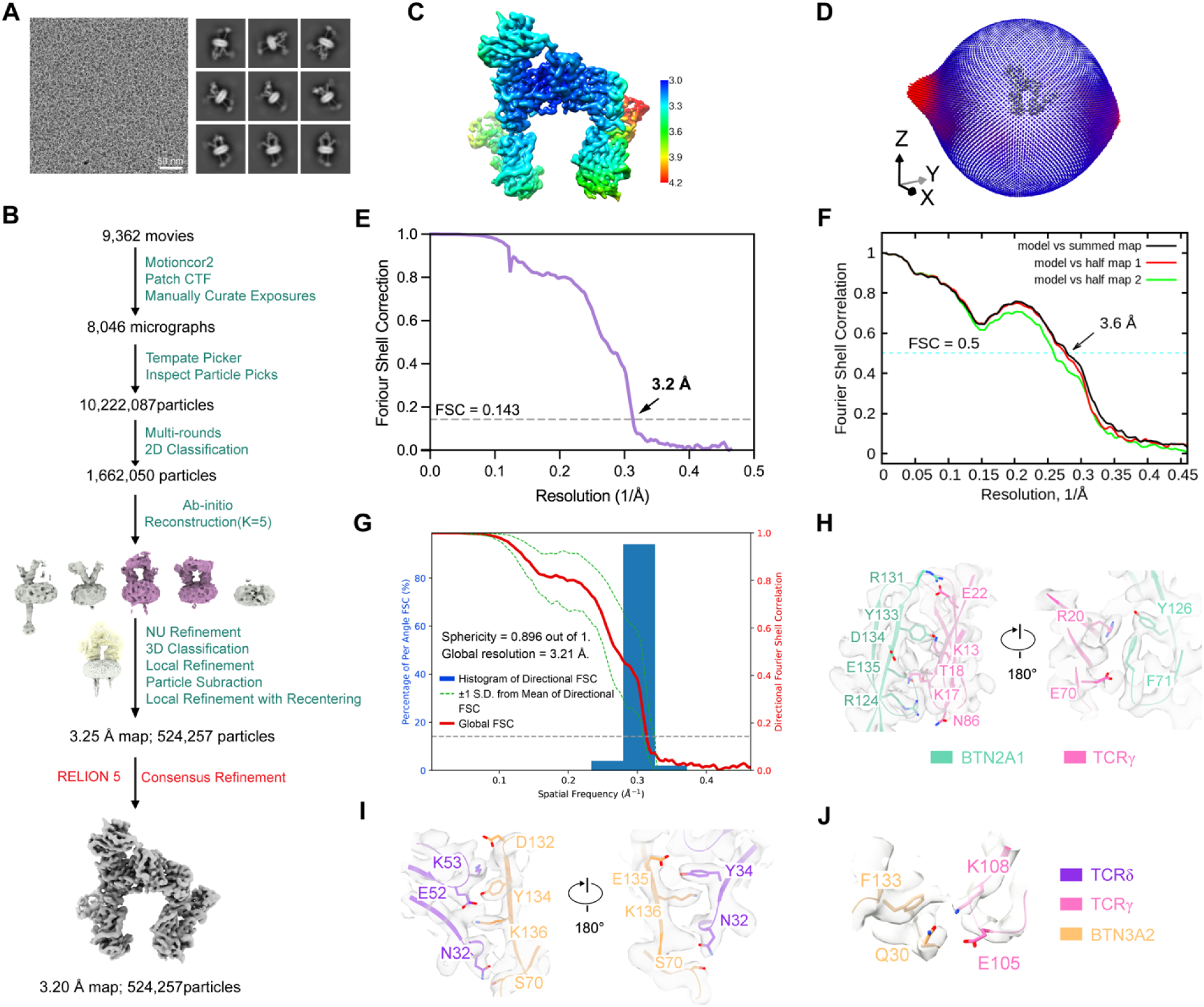
Cryo-EM data processing and analysis of the G115 TCR–AAR complex. (A and B) Representative cryo-EM images, 2D class averages, and data processing workflows for the G115 TCR–AAR complex. AAR complex, a 3A2 complex variant with mutations H381R in BTN3A1 and R477A/T510A in BTN2A1 B30.2 domains to disrupt pAg binding. (C and D) Local resolution estimation (C) and angular distribution (D) for the final reconstructions. The map is contoured at 9σ in (C). (E) Fourier shell correlation (FSC) curves for the final refinement. The dashed line indicates FSC = 0.143. (F) Model versus map FSC curves for the G115 TCR–AAR complex. (G) 3DFSC plots for the G115 TCR–AAR complex. (H–J) Close-up views of the interfaces between IgV_BTN2A1_ and the Vγ9 domain (H), IgV_BTN3A2_ and the Vδ2 domain (I), and IgV_BTN3A2_ and the Vγ9 domain (J) for the G115 TCR–AAR complex. The map is contoured at 9σ.

**Figure S15.**
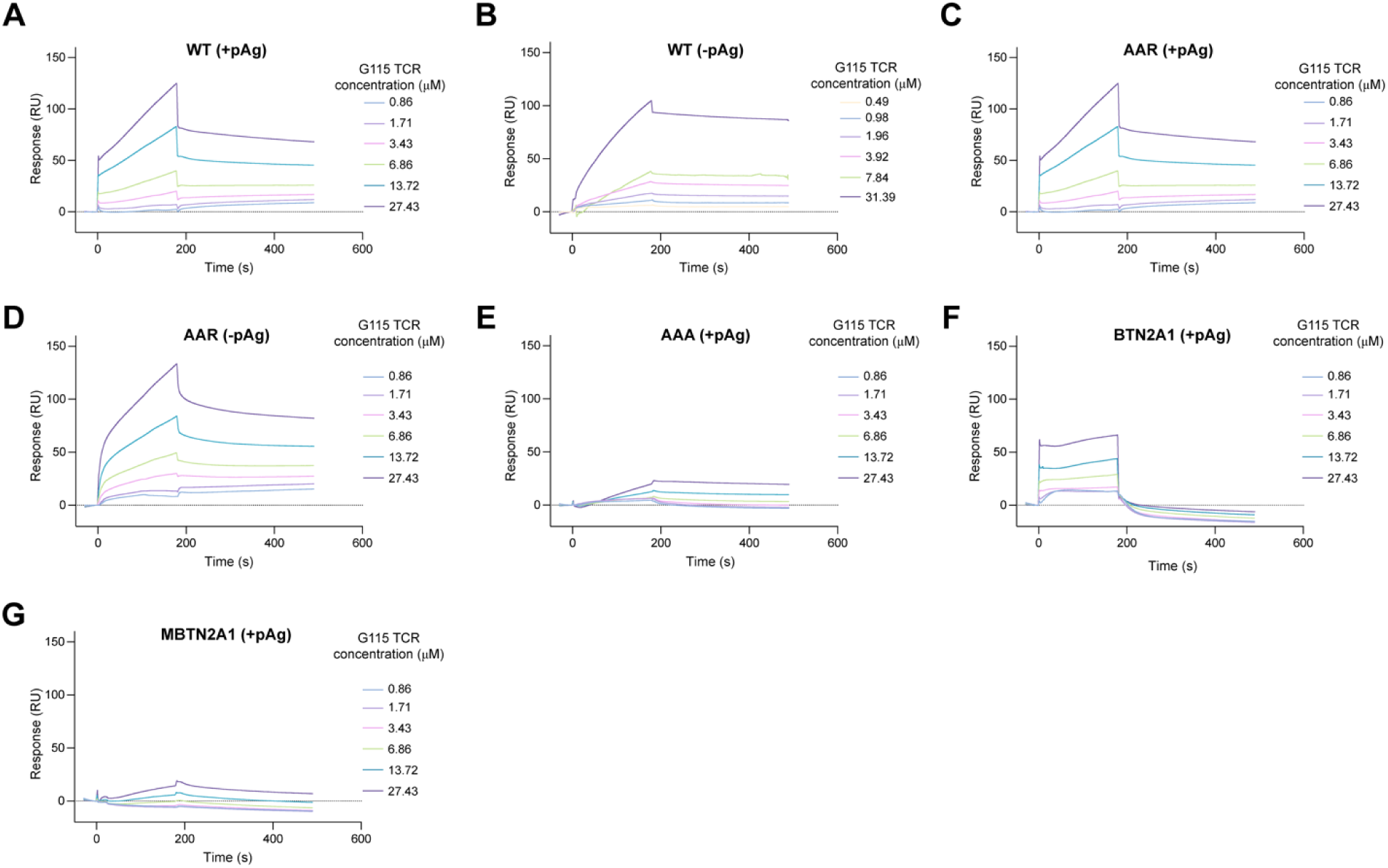
Surface plasmon resonance (SPR) analysis of binding affinities of G115 TCR towards different BTN molecules. (A–G) Sensorgrams of SPR showing the binding of soluble G115 TCR ectodomain to immobilized different BTN molecules. The presence or absence of HMBPP is indicated as +pAg or −pAg, respectively. WT: wild-type. AAR complex, a 3A2 complex variant with mutations H381R in BTN3A1 and R477A/T510A in BTN2A1 B30.2 domains, designed to disrupt HMBPP binding. AAA complex, a 3A2 complex variant carrying the R124A/Y133A mutations in BTN2A1 and D132A in BTN3A2. MBTN2A1, a BTN2A1 variant carrying E63A/R65A/F71A/K79A/R84A/E87A/R131A/E135A mutations in BTN2A1. RU: resonance unit.

**Figure S16.**
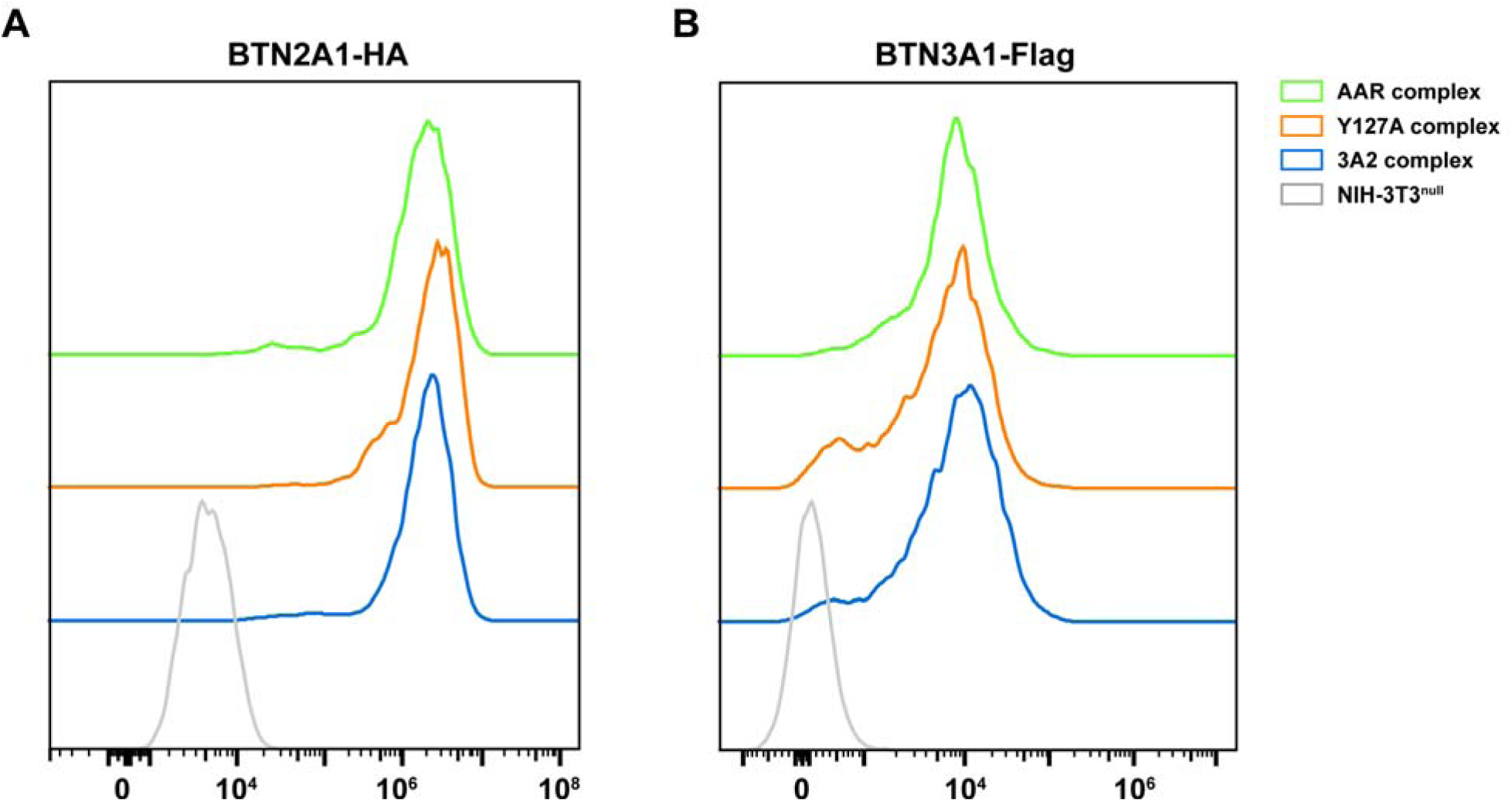
Surface expression of BTN2A1 and BTN3A in transduced NIH-3T3 cells expressing the WT 3A2 complex or its variants. (A) Anti-HA-BTN2A1 staining of each cell line with untransduced NIH-3T3 cells serving as negative controls (gray). (B) anti-Flag-BTN3A staining of each cell line with untransduced NIH-3T3 cells.

**Table S1.**
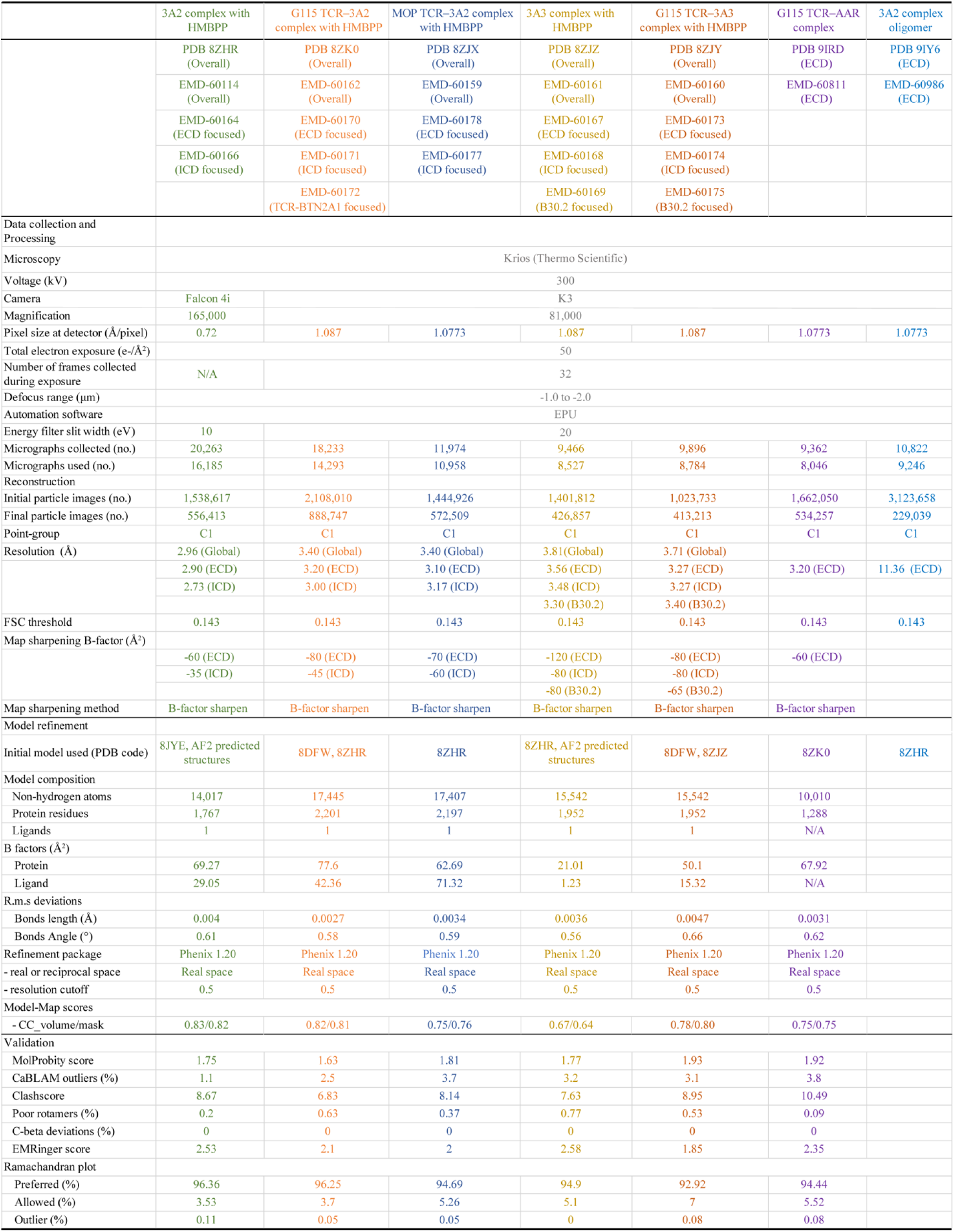
Cryo-EM data collection, refinement and validation statistics.

## References

1 Hayday, A., Dechanet-Merville, J., Rossjohn, J. & Silva-Santos, B Cancer immunotherapy by γδ T cells. Science 386, eabq7248. 10.1126/science.abq7248 (2024).

2 Vantourout, P. & Hayday, A Six-of-the-best: unique contributions of gammadelta T cells to immunology. Nat Rev Immunol 13, 88–100. 10.1038/nri3384 (2013).

3 Chien, Y. H., Meyer, C. & Bonneville, M. gammadelta T cells: first line of defense and beyond. Annu Rev Immunol 32, 121–155. 10.1146/annurev-immunol-032713-120216 (2014).

4 Davis, M. M. et al. Ligand recognition by αβ T cell receptors. Annual review of immunology 16, 523–544. (1998).

5 Mensurado, S., Blanco-Dominguez, R. & Silva-Santos, B. The emerging roles of gammadelta T cells in cancer immunotherapy. Nat Rev Clin Oncol 20, 178–191. 10.1038/s41571-022-00722-1 (2023).

6 Hayday, A. C. [gamma][delta] cells: a right time and a right place for a conserved third way of protection. Annu Rev Immunol 18, 975–1026. 10.1146/annurev.immunol.18.1.975 (2000).

7 Zheng, J., Liu, Y., Lau, Y. L. & Tu, W. gammadelta-T cells: an unpolished sword in human anti-infection immunity. Cell Mol Immunol 10, 50–57. 10.1038/cmi.2012.43 (2013).

8 Wu, Y. L., et al. gammadelta T cells and their potential for immunotherapy. Int J Biol Sci 10, 119–135. 10.7150/ijbs.7823 (2014).

9 Constant, P., et al. Stimulation of human gamma delta T cells by nonpeptidic mycobacterial ligands. Science 264, 267–270. 10.1126/science.8146660 (1994).

10 Tanaka, Y. et al. Natural and synthetic non-peptide antigens recognized by human gamma delta T cells. Nature 375, 155–158. 10.1038/375155a0 (1995).

11 Hintz, M. et al. Identification of (E)-4-hydroxy-3-methyl-but-2-enyl pyrophosphate as a major activator for human gammadelta T cells in Escherichia coli. FEBS Lett 509, 317–322. 10.1016/s0014-5793(01)03191-x (2001).

12 Zhao, L., Chang, W. C., Xiao, Y., Liu, H. W. & Liu, P. Methylerythritol phosphate pathway of isoprenoid biosynthesis. Annu Rev Biochem 82, 497–530. 10.1146/annurev-biochem-052010-100934 (2013).

13 Gober, H. J., et al. Human T cell receptor gammadelta cells recognize endogenous mevalonate metabolites in tumor cells. J Exp Med 197, 163–168. 10.1084/jem.20021500 (2003).

14 Benzaid, I. et al. High phosphoantigen levels in bisphosphonate-treated human breast tumors promote Vgamma9Vdelta2 T-cell chemotaxis and cytotoxicity in vivo. Cancer Res 71, 4562–4572. 10.1158/0008-5472.CAN-10-3862 (2011).

15 Ashihara, E. et al. Isopentenyl pyrophosphate secreted from Zoledronate-stimulated myeloma cells, activates the chemotaxis of gammadeltaT cells. Biochem Biophys Res Commun 463, 650–655. 10.1016/j.bbrc.2015.05.118 (2015).

16 Kunzmann, V., Bauer, E. & Wilhelm, M Gamma/delta T-cell stimulation by pamidronate. N Engl J Med 340, 737–738. 10.1056/nejm199903043400914 (1999).

17 Dieli, F. et al. Induction of gammadelta T-lymphocyte effector functions by bisphosphonate zoledronic acid in cancer patients in vivo. Blood 102, 2310–2311. 10.1182/blood-2003-05-1655 (2003).

18 Harly, C. et al. Key implication of CD277/butyrophilin-3 (BTN3A) in cellular stress sensing by a major human γδ T-cell subset. Blood 120, 2269–2279. 10.1182/blood-2012-05-430470 (2012).

19 Rigau, M., et al. Butyrophilin 2A1 is essential for phosphoantigen reactivity by gammadelta T cells. Science 367. 10.1126/science.aay5516 (2020).

20 Karunakaran, M. M. et al. Butyrophilin-2A1 Directly Binds Germline-Encoded Regions of the Vgamma9Vdelta2 TCR and Is Essential for Phosphoantigen Sensing. Immunity 52, 487–498 e486. 10.1016/j.immuni.2020.02.014 (2020).

21 Cano, C. E. et al. BTN2A1, an immune checkpoint targeting Vgamma9Vdelta2 T cell cytotoxicity against malignant cells. Cell Rep 36, 109359. 10.1016/j.celrep.2021.109359 (2021).

22 Karunakaran, M. M. et al. A distinct topology of BTN3A IgV and B30.2 domains controlled by juxtamembrane regions favors optimal human gammadelta T cell phosphoantigen sensing. Nat Commun 14, 7617. 10.1038/s41467-023-41938-8 (2023).

23 Vantourout, P. et al. Heteromeric interactions regulate butyrophilin (BTN) and BTN-like molecules governing gammadelta T cell biology. Proc Natl Acad Sci U S A 115, 1039–1044. 10.1073/pnas.1701237115 (2018).

24 Rhodes, D. A. et al. Activation of human gammadelta T cells by cytosolic interactions of BTN3A1 with soluble phosphoantigens and the cytoskeletal adaptor periplakin. J Immunol 194, 2390–2398. 10.4049/jimmunol.1401064 (2015).

25 Fichtner, A. S. et al. Alpaca (Vicugna pacos), the first nonprimate species with a phosphoantigen-reactive Vgamma9Vdelta2 T cell subset. Proc Natl Acad Sci U S A 117, 6697–6707. 10.1073/pnas.1909474117 (2020).

26 Compte, E., Pontarotti, P., Collette, Y., Lopez, M. & Olive, D Frontline: Characterization of BT3 molecules belonging to the B7 family expressed on immune cells. Eur J Immunol 34, 2089–2099. 10.1002/eji.200425227 (2004).

27 Rhodes, D. A., Reith, W. & Trowsdale, J Regulation of Immunity by Butyrophilins. Annu Rev Immunol 34, 151–172. 10.1146/annurev-immunol-041015-055435 (2016).

28 Zhu, Y. et al. Phosphoantigen-induced inside-out stabilization of butyrophilin receptor complexes drives dimerization-dependent γδ TCR activation. Immunity 58, 1646–1659.e1645. 10.1016/j.immuni.2025.04.012 (2025).

29 Zhang, M. et al. Structures of butyrophilin multimers reveal a plier-like mechanism for Vγ9Vδ2 T cell receptor activation. Immunity 58, 1660–1669.e1667. 10.1016/j.immuni.2025.05.011 (2025).

30 Su, Q., et al. Cryo-EM structure of the human IgM B cell receptor. Science 377, 875–880. 10.1126/science.abo3923 (2022).

31 Xin, W. et al. Structures of human gammadelta T cell receptor-CD3 complex. Nature 630, 222–229. 10.1038/s41586-024-07439-4 (2024).

32 Chen, M., Su, Q. & Shi, Y Molecular mechanism of IgE-mediated FcεRI activation. Nature 637, 453–460. 10.1038/s41586-024-08229-8 (2025).

33 Fulford, T. S. et al. Vgamma9Vdelta2 T cells recognize butyrophilin 2A1 and 3A1 heteromers. Nat Immunol 25, 1355–1366. 10.1038/s41590-024-01892-z (2024).

34 Yuan, L. et al. Phosphoantigens glue butyrophilin 3A1 and 2A1 to activate Vgamma9Vdelta2 T cells. Nature 621, 840–848. 10.1038/s41586-023-06525-3 (2023).

35 Kreiss, M. et al. Contrasting contributions of complementarity-determining region 2 and hypervariable region 4 of rat BV8S2+ (Vbeta8.2) TCR to the recognition of myelin basic protein and different types of bacterial superantigens. Int Immunol 16, 655–663. 10.1093/intimm/dxh068 (2004).

36 Pyz, E. et al. The complementarity determining region 2 of BV8S2 (V beta 8.2) contributes to antigen recognition by rat invariant NKT cell TCR. J Immunol 176, 7447–7455. 10.4049/jimmunol.176.12.7447 (2006).

37 Wang, H., Fang, Z. & Morita, C. T Vgamma2Vdelta2 T Cell Receptor recognition of prenyl pyrophosphates is dependent on all CDRs. J Immunol 184, 6209–6222. 10.4049/jimmunol.1000231 (2010).

38 Willcox, C. R. et al. Butyrophilin-like 3 Directly Binds a Human Vgamma4(+) T Cell Receptor Using a Modality Distinct from Clonally-Restricted Antigen. Immunity 51, 813–825 e814. 10.1016/j.immuni.2019.09.006 (2019).

39 Willcox, C. R. et al. Phosphoantigen sensing combines TCR-dependent recognition of the BTN3A IgV domain and germline interaction with BTN2A1. Cell Rep 42, 112321. 10.1016/j.celrep.2023.112321 (2023).

40 Yamashita, S., Tanaka, Y., Harazaki, M., Mikami, B. & Minato, N Recognition mechanism of non-peptide antigens by human gammadelta T cells. Int Immunol 15, 1301–1307. 10.1093/intimm/dxg129 (2003).

41 Sandstrom, A. et al. The intracellular B30.2 domain of butyrophilin 3A1 binds phosphoantigens to mediate activation of human Vgamma9Vdelta2 T cells. Immunity 40, 490–500. 10.1016/j.immuni.2014.03.003 (2014).

42 Gu, S. et al. Phosphoantigen-induced conformational change of butyrophilin 3A1 (BTN3A1) and its implication on Vγ9Vδ2 T cell activation. Proc Natl Acad Sci U S A 114, E7311–e7320. 10.1073/pnas.1707547114 (2017).

43 Haryadi, R. et al. Optimization of heavy chain and light chain signal peptides for high level expression of therapeutic antibodies in CHO cells. PLoS One 10, e0116878. 10.1371/journal.pone.0116878 (2015).

44 Mason, J. M., Hagemann, U. B. & Arndt, K. M Improved stability of the Jun-Fos Activator Protein-1 coiled coil motif: A stopped-flow circular dichroism kinetic analysis. J Biol Chem 282, 23015–23024. 10.1074/jbc.M701828200 (2007).

45 Su, Q. et al. Structural basis for Ca(2+) activation of the heteromeric PKD1L3/PKD2L1 channel. Nat Commun 12, 4871. 10.1038/s41467-021-25216-z (2021).

46 Su, Q. et al. Cryo-EM structure of the polycystic kidney disease-like channel PKD2L1. Nat Commun 9, 1192. 10.1038/s41467-018-03606-0 (2018).

47 Su, Q., et al. Structure of the human PKD1-PKD2 complex. Science 361. 10.1126/science.aat9819 (2018).

48 Lei, J. & Frank, J. Automated acquisition of cryo-electron micrographs for single particle reconstruction on an FEI Tecnai electron microscope. J Struct Biol 150, 69–80. 10.1016/j.jsb.2005.01.002 (2005).

49 Zheng, S. Q., et al. MotionCor2: anisotropic correction of beam-induced motion for improved cryo-electron microscopy. Nat Methods 14, 331–332. 10.1038/nmeth.4193 (2017).

50 Punjani, A., Rubinstein, J. L., Fleet, D. J. & Brubaker, M. A. cryoSPARC: algorithms for rapid unsupervised cryo-EM structure determination. Nat Methods 14, 290–296. 10.1038/nmeth.4169 (2017).

51 Punjani, A., Zhang, H. & Fleet, D. J Non-uniform refinement: adaptive regularization improves single-particle cryo-EM reconstruction. Nat Methods 17, 1214–1221. 10.1038/s41592-020-00990-8 (2020).

52 Burt, A. et al. An image processing pipeline for electron cryo-tomography in RELION-5. FEBS Open Bio 14, 1788–1804. 10.1002/2211-5463.13873 (2024).

53 Asarnow, D., Palovcak, E. & Cheng, Y UCSF pyem v0. 5. Zenodo 3576630. 10.5281/zenodo (2019).

54 Zivanov, J., Nakane, T. & Scheres, S. H. W A Bayesian approach to beam-induced motion correction in cryo-EM single-particle analysis. IUCrJ 6, 5–17. 10.1107/s205225251801463x (2019).

55 Tan, Y. Z., et al. Addressing preferred specimen orientation in single-particle cryo-EM through tilting. Nature methods 14, 793–796. 10.1038/nmeth.4347 (2017).

56 Meng, E. C. et al. UCSF ChimeraX: Tools for structure building and analysis. Protein Sci 32, e4792. 10.1002/pro.4792 (2023).

57 Pettersen, E. F. et al. UCSF Chimera—a visualization system for exploratory research and analysis. Journal of computational chemistry 25, 1605–1612. 10.1002/jcc.20084 (2004).

58 Fulford, T. S. et al. Vγ9Vδ2 T cells recognize butyrophilin 2A1 and 3A1 heteromers. bioRxiv, 2023.2008.2030.555639. 10.1101/2023.08.30.555639 (2023).

59 Jumper, J. et al. Highly accurate protein structure prediction with AlphaFold. Nature 596, 583–589. 10.1038/s41586-021-03819-2 (2021).

60 Liebschner, D. et al. Macromolecular structure determination using X-rays, neutrons and electrons: recent developments in Phenix. Acta Crystallogr D Struct Biol 75, 861–877. 10.1107/s2059798319011471 (2019).

61 Emsley, P., Lohkamp, B., Scott, W. G. & Cowtan, K. Features and development of Coot. Acta Crystallogr D Biol Crystallogr 66, 486–501. 10.1107/s0907444910007493 (2010).

62 Tang, G., et al. EMAN2: an extensible image processing suite for electron microscopy. J Struct Biol 157, 38–46. 10.1016/j.jsb.2006.05.009 (2007).

63 Williams, C. J., et al. MolProbity: More and better reference data for improved all-atom structure validation. Protein Sci 27, 293–315. 10.1002/pro.3330 (2018).

64 Costantini, L. M. et al. A palette of fluorescent proteins optimized for diverse cellular environments. Nat Commun 6, 7670. 10.1038/ncomms8670 (2015).

65 Lee, J. et al. Versatile phenotype-activated cell sorting. Sci Adv 6. 10.1126/sciadv.abb7438 (2020).

